# Wunen(s) help navigate Primordial Germ Cells by attenuating Hedgehog signaling

**DOI:** 10.64898/2026.05.01.722161

**Authors:** Amrita Roy, Adheena Elsa Roy, Airat Ibragimov, Juliana DaSilva, Kundan Kumar, Paul Schedl, Siddhesh S Kamat, Girish S Ratnaparkhi, Girish Deshpande

## Abstract

Embryonic PGC migration in *Drosophila* is orchestrated by non-cell autonomous repulsive and attractive cues, controlled by Wunen(s) and, HMGCoA-reductase (Hmgcr), respectively. Hedgehog (Hh), a PGC attractant, is potentiated by Hmgcr. We demonstrate that Wunen(s) employ both nonautonomous and autonomous modes to inhibit Hh signaling. In embryos maternally compromised for *wunen*, mesodermal cells and PGCs accumulate Hh, leading to precocious clumping of the PGCs, also observed upon PGC-specific loss of *patched* (*ptc*) – the Hh receptor and an antagonist of Smoothened (Smo), a G protein-coupled receptor (GPCR), involved in Hh signal transduction. Consistently, Wunen(s) inhibit membrane localization of Smo. Conversely, simultaneous overexpression of *wunen* mitigates PGC scattering induced by ectopic *hmgcr* expression. Remarkably, lipid metabolism is involved both in fine-tuning Hh signalling and choreographing PGC migration. Lipidomics of embryonic extracts after maternal knockdown of *wunen* confirms qualitative and quantitative disruptions in lipid composition. Here, we discuss the ability of Wunen(s) to repress Hh signalling vis-a-vis their repellent activity during PGC migration.

## Introduction

In diverse developmental contexts, directed cell migration constitutes a conserved and vital feature underlying the formation of a variety of cellular assemblies, including tissues and organs (Araujo, 2015; Campanale and Montell, 2023; Chen et al., 2025; Chowdhury et al., 2026; Diaz et al., 2025; Roper, 2025; Schick et al., 2025). Migrating cells navigate across multiple cell layers and travel long distances, which is controlled by soluble factors, chemokines, and intercellular communication, including cell adhesion (Barton et al., 2024; Paluch et al., 2016). Intriguingly, at times, proper migration of a specific cell type is contingent upon secreted signaling ligands functioning either as attractive or repulsive cues. Because these migratory events are essential for generating fully functional and fertile multicellular organisms, a wide range of model systems, spanning from *Dictyostelium* to higher vertebrates, including mammals, are used to elucidate the underlying mechanisms (Deshpande et al., 2025a; Santos and Lehmann, 2004a). Given the biological relevance of these processes, severe cell migration aberrations have detrimental consequences, including sterility, metastasis and lethality (Barton et al., 2024; Chowdhury et al., 2026; Paluch et al., 2016; Roper, 2025).

Embryonic gonad coalescence in *Drosophila* has emerged as a formidable context to analyze the mechanistic underpinnings of cell migration (Chen et al., 2025; Deshpande et al., 2025a; Kunwar et al., 2006). Several features render it an ideal system for uncovering and elucidating multimodal mechanisms underlying cell migration. First, the embryonic *Drosophila* gonad consists of just two cell types: Primordial Germ Cells (PGCs) and Somatic gonadal precursor cells (SGPs), which are specified at distant locations, with the PGCs undergoing directed migration towards the SGPs. PGCs are initially formed at the posterior of the embryo and, at the onset of gastrulation, are located next to a layer of somatic cells. During gastrulation they are internalized by the invagination of the midgut. At the germband extension (stage 9), PGCs localized at the blind end of the gut begin to segregate away from one another and individualize in a coordinated manner before traversing through the gut epithelium. After emerging from the gut around stage 10, the migrating PGCs split into two bilateral clusters, enter the mesoderm and begin migrating towards the SGPs. The SGPs are specified in para-segments 10-13 and are arranged in two bilateral clusters. Once the PGCs and SGPs come into contact, they intermingle and eventually compact into a tight cluster to form the paired primitive embryonic gonad.

The migration of the PGCs from the blind end of the gut and through the mesoderm is orchestrated by guidance signals emanating from the SGPs (Barton et al., 2024; Chen et al., 2025; Deshpande et al., 2025b). The first hint of the possibility came from the discovery that 3-hydroxy-3-methylglutaryl-CoA reductase (Hmgcr) is responsible for the production or transmission of an attractive guidance cue. *hmgcr* is the rate-limiting step in several biosynthetic pathways including that of cholesterol and isoprenoids. Loss of *hmgcr* results in the scattering or mis-migration of PGCs in the posterior of the embryo, while ectopic expression of *hmgcr* induces PGCs to migrate abnormally towards the cells/tissues expressing *hmgcr*. In addition, *hmgcr* is initially expressed broadly in the mesoderm; however, as the embryo develops, its expression becomes progressively restricted, first to the posterior mesoderm and by stage 12, almost exclusively in the SGPs (Barton et al., 2024).

Though these studies demonstrated that *hmgcr* plays a key role in orchestrating PGCs migration, its immediate biosynthetic product (mevalonate) does not appear to be a plausible candidate as a guidance cue for the migrating PGCs (Santos and Lehmann, 2004b). However, since mis-expression of *hmgcr* in many different cell types/tissues in the embryo induced PGCs to migrate towards the site of *hmgcr* expression, it was apparent that the actual attractant must be broadly expressed in the embryo but somehow limited in its ability to function as an attractant in the absence of *hmgcr*. The identification of Hedgehog (Hh) as the likely PGC attractant (Deshpande et al., 2001) supported this notion. Like *hmgcr*, Hh is expressed in the SGPs and their progenitors. Likewise, misexpression of *hh* induces a subset of PGCs to migrate towards the ectopic source of Hh. Interestingly, the mis-migration of PGCs induced by *hh* ectopic expression is typically less severe than that induced by *hmgcr* expression in the same cell types/tissues (Deshpande et al., 2013). This observation suggested that *hmgcr* was a limiting factor in Hh ligand transmission. Several findings support this idea. First, in the ectoderm of *hmgcr* mutant embryos, the Hh ligand is retained in *hh*-expressing cells instead of being transmitted to the neighboring cells (Deshpande and Schedl, 2005). Second, co-ectopic expression of *hh* and *hmgcr* in the CNS leads not only to much more severe PGC mis-migration, but also substantially enhances the transmission of the Hh ligand from the ectoderm into the underlying mesoderm where it decorates the migrating PGCs (Deshpande et al., 2013). Third, epistasis experiments show that *hh* functions downstream of *hmgcr* in PGC migration: the effects of ectopic *hmgcr* expression in the CNS on PGC migration can be suppressed by simultaneous knockdown of *hh* (Deshpande et al., 2023). Fourth, two other genes, *toutvelu* and *dispatched*, known to be important for the cytoneme-based transmission of the Hh ligand are also epistatic to ectopically expressed *hmgcr* (Deshpande et al., 2023).

Long-distance transmission of the Hh ligand requires proteolytic cleavage of the precursor protein and concurrent cholesterol modification. While several studies indicate that cholesterol modification is important for the transmission of the Hh ligand from the SGPs to the migrating PGCs (Deshpande et al., 2016), flies lack the enzymes downstream of *hmgcr* for biosynthesis of cholesterol and thus this cannot be the function of *hmgcr* in the fly Hh signalling pathway. A solution to this puzzle came from the studies of (Santos and Lehmann, 2004b), who showed that genes in the isoprenoid branch of the *hmgcr* biosynthetic pathway (Farnesyl diphosphate synthase: *fpps*; Geranyl-geranyl diphosphate synthase: *qme*) function downstream of *hmgcr* in PGC migration as does the isoprenoid transferase Geranyl-geranyl transferase (GGT1). The isoprenoid biosynthetic pathway is also required for the release/transmission of the Hh ligand, and at least one target in the Hh signaling pathway for geranylation by GGT1 is the gγ subunit of the heterotrimeric G protein complex (Deshpande et al., 2009).

Further evidence that the Hh signaling pathway orchestrates PGCs migration came from the experiments demonstrating that the reciprocal activities of the two receptors of the Hh ligand, namely Patched (Ptc) and Smoothened (Smo). Germline line clones and ectopic expression experiments show that the Hh receptor Patched (Ptc) is required in the PGCs for proper migration (Deshpande et al., 2001). When *ptc* function is compromised in PGCs, they tend to clump together as would be expected if they constitutively “receive” a signal. Reception of the Hh ligand by the PGCs results in the internalization of Hh and Ptc into large intercellular puncta (Kim et al., 2021) whereas the Smoothened (Smo) protein associates with the membrane in an actin-rich protrusion at the leading edge of the migrating PGCs (Kim et al., 2021). Contrary to Ptc, when Smo activity is disrupted, the PGCs scatter in the mesoderm (Deshpande et al., 2001). The non-canonical target for Smo activation is a G-protein coupled receptor, Trapped in Endoderm (Tre1). Like Smo, Tre1 localizes to the protrusions at the migration front together with dPIP5k (phosphatidylinositol 4-phosphate 5-kinase), the enzyme responsible for generating PI(4, 5)P_2_ (Kim et al., 2021). PIP5k targeted to the migration front by Tre1 induces the local accumulation of PI(4, 5)P_2_ which, via Wiskott–Aldrich syndrome protein (WASP), activates Arp2/3 dependent actin polarization at the leading edge. In *tre1* mutants PIP5K does not localize to the migration front and as a result the production of PI(4, 5)P_2_ does not become polarized. Likewise, actin is not preferentially polymerized at the migration front, thus linking the downstream events to the initial targeting of Smo to the membrane at the migration front in response to the reception of the Hh ligand (Kim et al., 2021).

While Hh expressed by the SGPs induces PGC migration towards the SGPs by functioning as an attractant, there are two related lipid phosphate phosphatases (LPPs) *wunen* (*wun*) and *wunen-2* (*wun2*) that guide PGC migration by generating, in some as yet unknown manner, the repulsive signals (Hanyu-Nakamura et al., 2004; Renault and Lehmann, 2006; Starz-Gaiano et al., 2001). Both *wun* and *wun2* are maternally deposited, and the maternal products are found in both the soma and the PGCs. They are also zygotically expressed in the epidermis, gut primordium, and nervous system. Intriguingly, *wun* or *wun2* transcripts are missing from the blind end of the gut primordium, a site that coincides with the region where PGCs undergo transepithelial migration across the gut. Both ‘loss’ and ‘gain’ of function experiments have indicated that Wun(s) in the soma (of maternal and zygotic origin) repel migrating PGCs, and that PGCs avoid invading the ‘wrong’ tissues because of Wun(s). For instance, PGCs bypass the nervous system due to endogenous expression of *wun*(s) in the ventral nerve chord (Hanyu-Nakamura et al., 2004; LeBlanc and Lehmann, 2017; Renault et al., 2010; Renault et al., 2004; Zhang et al., 1997). Conversely, when either *wun* or *wun2* is expressed in the mesoderm, PGCs are unable to enter the mesoderm. In addition to this non-autonomous repulsive function, the Wun(s) are also required in migrating PGCs where they contribute to their individualization i.e. PGC-PGC repulsion and help facilitate their proper migration (Renault et al., 2010; Renault et al., 2004).

The *wun(s*) have been studied extensively since their discovery in the ‘90s (Hanyu-Nakamura et al., 2004; LeBlanc and Lehmann, 2017; Renault et al., 2010; Renault et al., 2004; Zhang et al., 1997). However, the biochemical and genetic characterisation of Wunen(s) as lipid phosphate phosphatases has not uncovered any specific molecular substrate(s). Thus, the mechanistic basis for their repulsive activity in the soma and their cell-autonomous function(s) in PGCs have remained a mystery. One candidate target that could potentially connect both their non-autonomous function in the soma and their cell-autonomous function in PGCs is the Hh signalling pathway. This is especially relevant as Hh pathway components, including Hh itself, undergo lipid modifications which regulate range and/or strength of Hh signaling in different developmental contexts. Here we show that *wun(s*) suppress Hh signalling in both the soma and migrating PGCs. In the soma, their activity counteracts *hmgcr*-dependent potentiation of Hh transmission. In PGCs, *wun(s)* appear to suppress the response to Hh ligand reception. In addition, we demonstrate that, Suppressor of actin 1 (Sac1), a lipid phosphatase, plays a similar role in PGC migration by downregulating Hh signalling. To better understand the activities of the two enzymes involved in lipid metabolism, Wun(s) and Sac1, we compared unbiased lipidomes from the respective embryonic extracts. Intriguingly, these lipid metabolism enzymes appear to exert their effects on the Hh signaling pathway by targeting different phospholipids (Burnett and Howard, 2003; Burnett et al., 2004). Taken together, our data underscore the complementary influence of both attractive and repulsive arms on the same guidance cue. This balance may prove to be highly informative in other developmental contexts including normal as well as pathological conditions.

## Results

### Embryos derived from mothers compromised for *wun* display enhanced levels of Hh targets

In stage 8-11 embryos, a positive autoregulatory loop exists in the embryonic ectoderm in which *hh* activates *wingless* (*wg*) expression in Hh-receiving cells, and *wg* in turn upregulates *engrailed* (*en*) expression in *hh*-expressing cells (Fig. 1A-B). We previously showed that mutations in *hmgcr* disrupt this autoregulatory loop, leading to significant downregulation of En expression. Thus, if the Wun(s) function to suppress the *hh* signaling pathway, we might expect to observe an opposite phenotype—namely that En protein expression will be upregulated when their activity is compromised.

**Figure 1.**
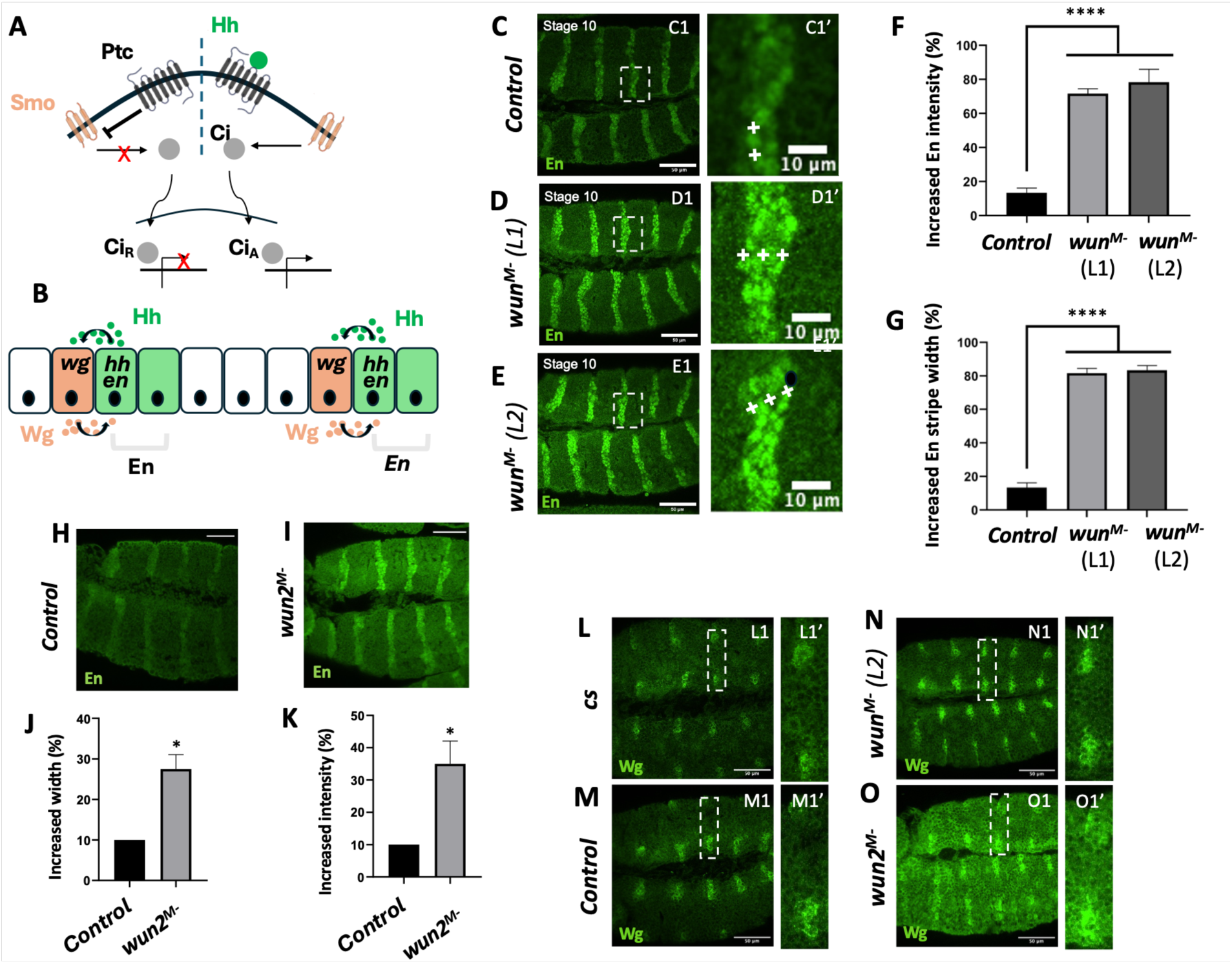
Embryos either maternally (*M^-^*) or zygotically (*Z^-^*) compromised for *wun* display enhanced levels of ectodermal En and Ptc. **(A-B)** Schematics depict the canonical Hh signalling pathway (A), where in the ‘signaling Off/Hh-unbound’ state, Ptc inhibits Smo, and in the absence of signal transduction downstream of Smo, Ci retains its repressor activity (CiR). In the ‘On’ state, Hh is bound to its receptor Ptc, which in turn relieves Smo inhibition. Smo dependent signaling results in generation of transcriptional activator (CiA) form. Panel (B) represents Hh’s role in the formation of segmental boundaries in the embryonic epidermis. En, expressed in the posterior of each para-segment, activates *hh.* Hh is secreted from the En-positive cells, and binds to Ptc receptor in the neighboring cells, wherein activation of Hh signaling leads to *wg* expression. Wg, in turn, is secreted from these neighboring cells, and signals reciprocally to the En-positive cells, setting up a tri-partite positive feedback loop between En/Wg/Hh. Together the activities of these three gene products determine the para-segmental boundary and thus defining the segment polarity. **(C-E)** *nos-gal4* virgin females were mated with males of specified genotypes, including negative control (*UAS-lex RNAi*) and two different *UAS-wun RNAi* lines, denoted as *wun Line1(L1;* BDSC 32429) and *wun Line2 (L2;* BDSC 32430). These knockdowns are hypomorphic. Stage 10 embryos derived from mated females carrying both transgenes (*nos-gal4/UAS-RNAi*) were stained with anti-Engrailed (En) antibody. Compared to the *Control lex^M-^*embryos (A), a significantly higher percentage of *wun^M-^* embryos (B and C) display increased En protein levels and broader En stripes. The two *UAS-wun RNAi* lines behaved similarly. Panel A1’-C1’ shows magnified insets that demonstrate both the elevation in the En levels and broadening of the stripes, indicating additional number of nuclei that stain for En. The ‘+’ mark denotes En positive cells. **(F-G)** Quantification of percentage of embryos showing increased En intensity and broader stripes (n=15, N=3, ****p<0.0001). Significance was estimated using Ordinary one-way ANOVA. **(H-I)** Stage 10 embryos maternally depleted of control *lexA* and *wun2*. Compared to the negative control *lex^M-^* embryos (H), a higher percentage of *wun2^M-^* embryos display as broader En stripes and increased levels of En protein (I). (J-K) Quantification of En stripe broadening (J) and increased En expression (K) in embryos. Significance was estimated using one-way ANOVA (n=10 N=2), *p<0.05. **(L-O)** *nos-gal4* virgin females were mated with males of specified genotypes, including negative control (*UAS-lex RNAi,* M), *UAS-wun RNAi* (N) and *UAS-wun2 RNAi* (O). CS embryos (L) were an additional control. Stage 10 embryos derived from mated females carrying both transgenes (*nos-gal4/UAS-RNAi*) were stained with anti-Wingless (Wg) antibody. Compared to the Controls (L,M), a significantly higher percentage of *wun^M-^ & wun2^M-^* embryos (N and O, respectively) display an increased level of Wg protein. Panel L1’-O1’ shows magnified insets that demonstrate the elevation in Wg levels.

As a test of this idea, *wun* was *maternally* (M-) depleted, by driving the expression of *UAS-wunRNAi* in the adult female germline using *nos-Gal4* (Fig. 1 Fig. 1C-E). We then examined En protein expression in the progeny (F2) of these females (mated to wild-type males). At stage 10, En expression in these *wun^M-^*embryos is appreciably increased compared to wild-type (Fig.1, C1 versus D1, E1). In addition to a higher level of expression in individual cells, there is a readily discernible increase in the number of En-positive cells per stripe (Fig. 1, C1’ vs D1’, E1’). Upregulation of En expression together with a broadening of En stripe is also observed in *wun2^M-^*embryos (generated using the same strategy), although the phenotype is weaker compared to *wun^M-^* (Suppl. Fig.1). Thus, compromising the activity of either *wun* or *wun2* maternally is sufficient to upregulate Hh signaling in the ectoderm. Both *wun* and *wun2* are maternally and zygotically expressed. While considerably higher levels of *wun2* are maternally deposited, rescue experiments have confirmed that they are functionally analogous. It should be noted that compromising *wun* may yield more penetrant somatic defects whereas reducing *wun2* may result in more severe PGC-related phenotypes.

To confirm the somatic defects induced by the loss of *wun*, we also analyzed expression of Wingless (Wg), another target of Hh signaling in the embryonic ectoderm. Analogous to En staining pattern, Wg expression is also enhanced upon compromising Wun activity. (Of note, the increase in both En and Wg levels seems to be confined to the segments alone and the upregulation did not appear to be random).

In addition to being maternally deposited, the *wun(s)* are zygotically expressed in the gut endoderm and in a segmentally repeating pattern in the embryonic ectoderm. To determine whether zygotically expressed *wun* also functions to suppress Hh signaling, we mated females carrying a *maternal-α−tubulin Gal4* transgene (*Mat Gal4*) to *UAS-wunRNAi* males. Suppl. Fig. 2 shows that En expression is upregulated in these F1 embryos as compared to the control.

These findings indicate that both maternal and zygotic components of *wun* function to suppress *hh* signaling. If this is correct, then ectopic expression of *wun* in the zygote should suppress *hh* signaling and result in a decrease in En expression. To test this prediction, we mated females carrying *Mat Gal4* to males carrying a *UAS* transgene encoding a *wunen-GFP* fusion protein. As shown in Suppl. Fig. 2C, zygotic overexpression of *wun* (Wun^OE^) in the embryo decreases En protein expression.

### Embryos zygotically compromised for wun display increased levels of β-galactosidase driven by patched-Gal4

The interaction between Hh and Ptc results in the translocation of the Smo protein to the membrane and signal transduction (Hu and Song, 2019; Ingham, 2022; Zhang and Beachy, 2023) (HU AND SONG 2019; INGHAM 2022; ZHANG AND BEACHY 2023) (Fig. 1A, B). In the canonical pathway, this results in the transcriptional upregulation of *ptc* via Cubitus interruptus (Ci). Ptc upregulation provides a negative feedback loop that limits Hh activity and spread (Briscoe et al., 2001; Casali and Struhl, 2004; Chen and Struhl, 1998). If *wun* functions to suppress this branch of the canonical *hh* pathway, compromising *wun* activity zygotically (Z-) should result in a detectable increase in *ptc* transcriptional activation. We mated females carrying a *ptc-gal4 UAS-β-gal* (Johnson et al., 2000) recombinant to either control males or males carrying the *UAS-wun RNAi* transgene and stained the resulting (F1) embryos with anti-*β-gal* (anti-*β-*gal) antibodies. The control embryos displayed a segmental pattern that mimics *ptc* expression (Fig. 1H). By contrast, the *ptc-gal4 UAS-β-gal / UAS-wun RNAi* embryos (Fig. 1I-J) displayed a considerable increase in the anti-*β-*gal specific signal. These findings were subsequently confirmed by anti-Ptc antibody staining in embryos of the same genotype.

### Simultaneous overexpression of wun in the nervous system can mitigate PGC migration defects induced by elav-gal4 UAS-hmgcr

While the experiments involving RNAi-based embryonic knockdown suggested that *wun(s)* function to downregulate the *hh* signalling pathway in embryos, they do not provide insight into the underlying mechanism. Interestingly, similar to *wun(s),* another phospholipid phosphatase, *sac1,* functions to attenuate the *hh* signaling pathway (Yavari et al., 2010). Studies in wing and eye discs have argued that *sac1* downregulates the *hh* pathway, in the receiving cells, after reception of the Hh ligand. While a similar cell-autonomous mechanism downstream of reception could be deployed by the *wun(s)*, their ability to inhibit PGC migration into endodermal and ectodermal tissues suggests that they could also suppress the transmission of the Hh ligand. (The cell-autonomous and non-cell autonomous functions of Wun(s) need not be mutually exclusive). If this model is correct, then *wun(s)* would counteract the potentiation of Hh transmission by *hmgcr*.

Expression of *UAS-hmgcr* in the embryonic CNS using an *elav-Gal4* driver induces PGC migration defects. These PGC migration defects can be substantially enhanced by co-expressing *UAS-hh* in the CNS while they are suppressed by co-expressing *UAS-hhRNAi* (Deshpande et al., 2023). Likewise, *RNAi* knockdown in the CNS of two genes required for Hh transmission, *ttv* and *disp* also suppressed the PGC migration defects induced by ectopic *hmgcr (Deshpande et al., 2023)*. To explore the idea that *wun* functions to attenuate the transmission of the Hh ligand, we examined if overexpression of *wun* (*UAS-wun*) is also able to mitigate the PGC migration defects induced by ectopic expression of *hmgcr* in the embryonic nervous system using the *elav-Gal4* driver (embryos derived from the *elavGal4UAS-hmgcr* recombinant strain).

Stage 13-14 embryos were classified based on the number of mis-migrated/scattered PGCs: 0-4, 5-9, 10-14 and 15+ mis-migrated PGCs. Compared to control embryos (Fig. 2A and 2D), about 65% of *elav-gal4 UAS-hmgcr/+* embryo had 10 or more PGCs scattered/mis-migrated (Fig. 2B and 2E). As observed for *hh*, *ttv* and *disp* RNAi knockdowns, co-expression of Wun suppresses the PGC migration defects induced by *hmgcr*. Only about 20% of *elav-gal4 UAS-hmgcr/UAS-wunGFP* embryos had 10 or more mis-migrated PGCs (Fig. 2C and 2F). The suppression of mis-migration induced by *elav-gal4UAS-hmgcr* by simultaneous overexpression of *wun* is quantified in Fig. 2G. Taken together these data support the idea that PGCs are misdirected upon ectopic expression of *hmgcr* due to competition between the endogenous and ectopic sources of the attractant produced in the SGPs and the CNS respectively. This competition can be weakened by Wun overexpression in the CNS. The outcome is quantitatively similar to that of compromising positive regulators of Hh signaling such as *ttv* or *disp*.

**Figure 2.**
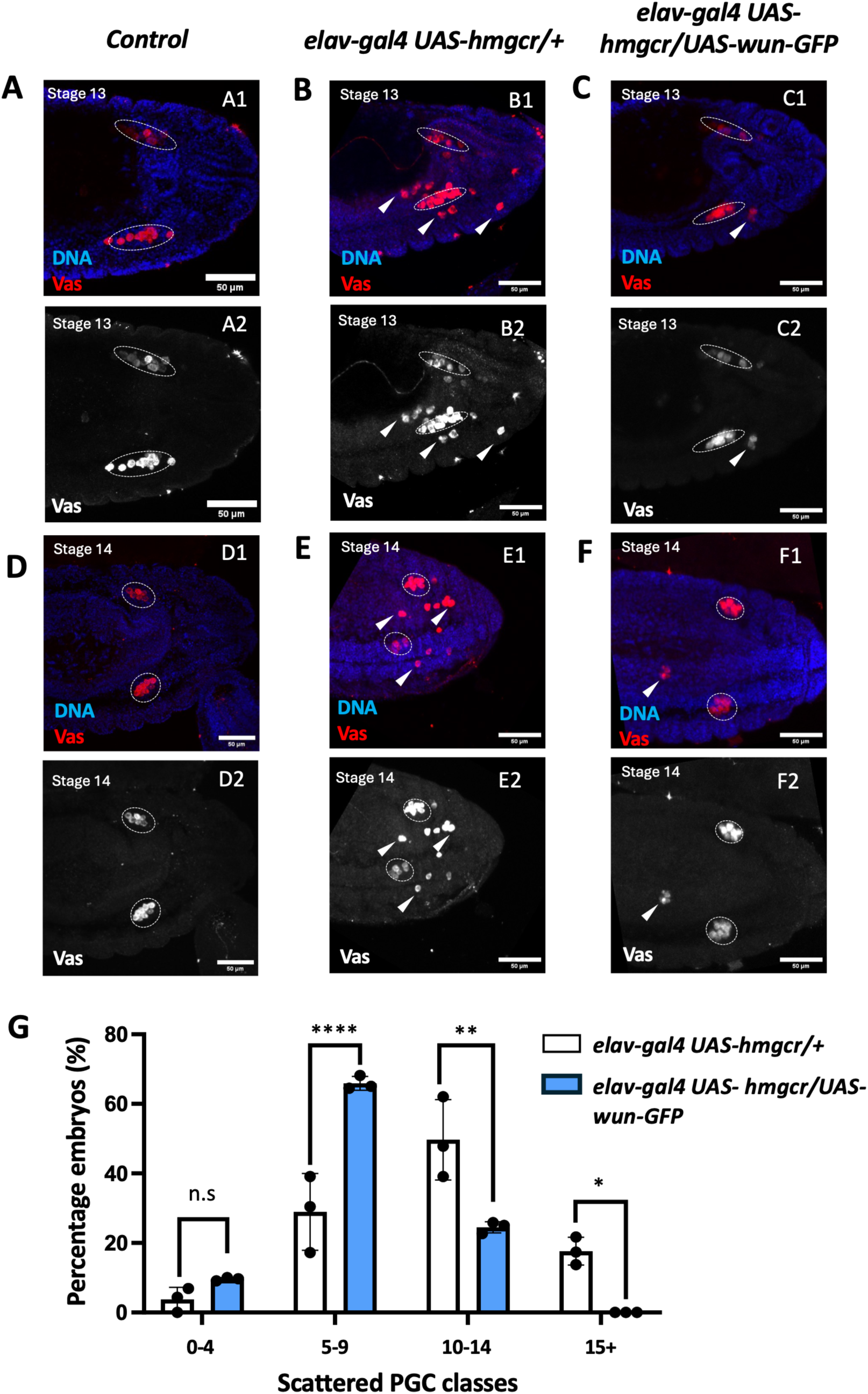
Aberrant PGC migration, induced by ectopic expression of hmgcr can be effectively suppressed by simultaneously overexpressing wun. **(A-C)** Stage 13 embryos. Compared to control CS (A) PGCs in elav-gal4 UAS-hmgcr/+ are greatly scattered away from the gonad (B). In elav-gal4 UAS-hmgcr/UAS-wun-GFP embryos, simultaneous overexpression of wun-GFP suppresses the migration defects induced by the ectopic expression of hmgcr in the nervous system (C) A2, B2, C2 are greyscale representations of the Vasa channel. **(D-F)** Stage 14 embryos of the indicated genotypes. Migration defects caused by ectopic expression of hmgcr in the nervous system are suppressed by simultaneous expression of wun-GFP. (A1-D1) Merged image with anti-Vas and counterstained with Hoechst. (A2-D2) PGCs stained with anti-Vas. D2, E2, F2 are greyscale representations of the vasa channel **(G)** Quantification of elav-gal4 UAS-hmgcr/+ and elav-gal4 UAS-hmgcr/UAS-wun-GFP embryos belonging to the mentioned classes of scattered PGCs (n=25, 3 replicates). Significance calculated using Two-way ANOVA and Tukey’s multiple comparison test ****p<0.0001, **p<0.01, *p<0.05.

### wun^M-^ embryos display elevated levels of endogenous Hh-GFP in the PGCs and surrounding mesodermal cells

When *hmgcr* is co-expressed with *hh-GFP* in the embryonic CNS using an *elav-Gal4* driver, it substantially enhances the transmission of the Hh-GFP ligand encoded by the transgene into and through the underlying mesoderm, where it decorates the migrating PGCs (Deshpande et al., 2023). Thus, a clear implication of the *hmgcr-wun* epistasis experiments in the previous section is that the *wun(s)* are able to negatively regulate the *hh* signaling pathway because they adversely influence the synthesis, stability, retention or efficient transmission of the Hh ligand. If this inference is correct, the pattern of accumulation of the endogenously expressed Hh should be substantially altered when *wun(s)* activity is compromised. To test this idea, we used the *nos-Gal 4*/*UAS-wun RNAi* (BDSC_32430) combination to deplete maternal deposition of *wun*. Females carrying this transgene combination were then mated to males carrying a BAC *hh-GFP* rescue construct (henceforth referred to as Hh-GFP), which recapitulates the expression pattern of the endogenous *hh* gene (Chen et al., 2017).

As shown in Fig. 3A-B, there is significant increase in the accumulation of the Hh ligand in in the somatic mesoderm of wun^M-^ embryos. As previously reported by (Kim et al., 2021) for wild type *hh-GFP* embryos, a substantial fraction of embryonic PGCs, display detectable levels of Hh-GFP from stage 9/10 onwards (Fig. 3D). Moreover, as predicted, there appeared to be a significant elevation in the number of Hh-GFP-puncta per PGC in the *wun^M-^* embryos (Fig. 3E-3F). To better analyze the change, we quantitated the total number of readily discernible Hh-GFP puncta per PGC in the control (*UAS-lexA RNAi*) and *wun^M-^* embryos. In control embryos, about 90% of the Hh-GFP signal-positive PGCs have 5 or less puncta and of these about 50% have only 1 or 2 Hh-GFP puncta. Since PGCs do not express Hh, the increase in Hh-GFP puncta/PGC in *wun^M-^* embryos must, at least in part, be due to the upregulation of Hh-GFP transmission from the surrounding mesodermal and ectodermal tissues. To confirm this idea, we measured the frequency of Hh-GFP puncta in a region (ROI) dorsal to the gut that is devoid of PGCs. As shown in Fig.3C, the number of Hh-GFP puncta in this region of the mesoderm is greatly enhanced in *wun^M-^* embryos. Taken together with the results in the previous sections, these findings indicate that *wun(s)* function to downregulate or block the transmission of Hh from cells that synthesize the ligand. This could also arise due to increased synthesis, stability of Hh ligand or its sequestration in a cellular compartment in the mesodermal cells. Future experiments will be necessary to delineate the precise mechanism underlying accumulation of excess Hh in the PGCs from *wun^M-^* embryos.

**Figure 3.**
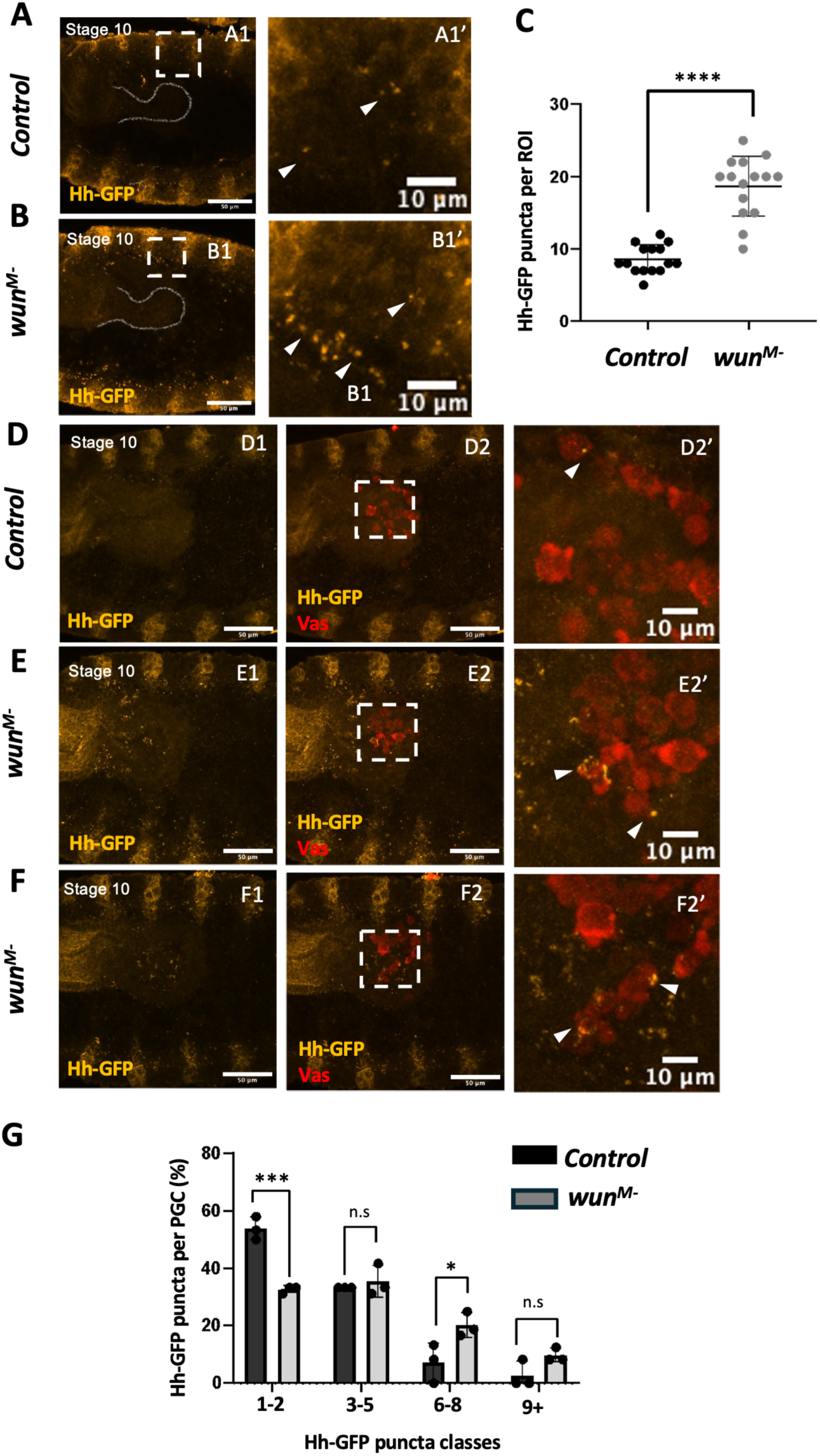
*wun^M-^ embryos* exhibit an increase in endogenous Hh-GFP puncta in the mesoderm and the adjacent PGCs. **(A-B)** Either *UAS-lexA RNAi* or *UAS-wun RNAi* males were mated with *nos-gal4*. virgin females carrying both the transgenes were mated with endogenous *Hh-GFP* males. Compared to control *lexA^M-^* (A), *wun^M-^* embryos show a higher number of *Hh-GFP* puncta (Yellow) in the mesoderm. **(C)** Individual Hh-GFP puncta were counted in 15 ROIs dorsal to the gut, per genotype (gut outline is marked in all the panels). The increase in Hh-GFP is quantified using an unpaired T-test, ****p<0.0001. (A1-B1). Magnified insets shown in (A1’-B1’), with arrowheads marking Hh-GFP puncta. **(D-F)** Compared to control, *lexA^M-^* (D) *wun^M-^* embryos show a higher number of Hh-GFP puncta in their PGCs (E and F), marked with anti-Vasa antibody (red) staining. Magnified insets (D2’-F2’), with arrowheads marking Hh-GFP puncta. **(G)** The total number of puncta was counted in 15 PGCS per genotype across three replicates, and the percentages of PGCs with 1-2, 3-5, 6-8 and 9+ Hh-GFP puncta are plotted. Significance was calculated using Two-way ANOVA and Tukey’s multiple comparison test (****p<0.0001, ***p<0.001, **p<0.01).

Furthermore, since the *wun(s)* are also required in PGCs (Renault et al., 2010; Renault et al., 2004), we cannot exclude a complementary cell autonomous effect in which the loss of maternally derived *wun* in migrating PGCs also contributes to the increase in PGC associated Hh-GFP puncta (see below).

### Aberrant localization of Smo in PGCs of wun^M-^ embryos

The reception of the Hh ligand results in the translocation of Smo from its cytoplasmic localization to the cell membrane. In PGCs Smo localizes to membrane protrusions at the leading edge of the migrating cells (Kim et al., 2021) and mediates the recruitment of the Tre1 GPCR receptor. Tre1 induces localized actin polymerization in response to a transduction pathway the produce PI(4,5)P_2_. Since Hh signaling to the migrating PGCs is polarized, there is typically only a single sharp focus of Smo protein which is localized at the leading edge (Fig. 4, A1–A4). Typically, wild-type PGCs display a single sharp focus with the residual protein being associated with the rest of the membrane. This pattern of localization is disrupted in *wun^M-^* (Fig. 4, B1-B4; C1–C4). To better understand the aberrant behavior, we classified individual PGCs based on the pattern of Smo localization. The classes included (i) PGCs showing no Smo-specific signal, (ii) PGCs showing a single focused crescent of Smo at the leading edge (iii) PGCs displaying diffuse localization of Smo across the membrane (iv) PGCs having a combination of a leading edge and a diffuse localisation of Smo (v) PGCs with multiple foci enriched for Smo-specific staining. In control PGCs, most PGCs have a single focus or a single focus surrounded by a more diffuse pattern of localization (∼60%). In contrast, less than 40% of the PGCs in *wun^M-^* embryos have a single focus or a single focus surrounded by a more pattern of localization. About 35% show strong staining across the entire cell, while over 10% have multiple Smo foci. The changes in Smo localization are quantitatively significant (Fig. 4D).

**Figure 4.**
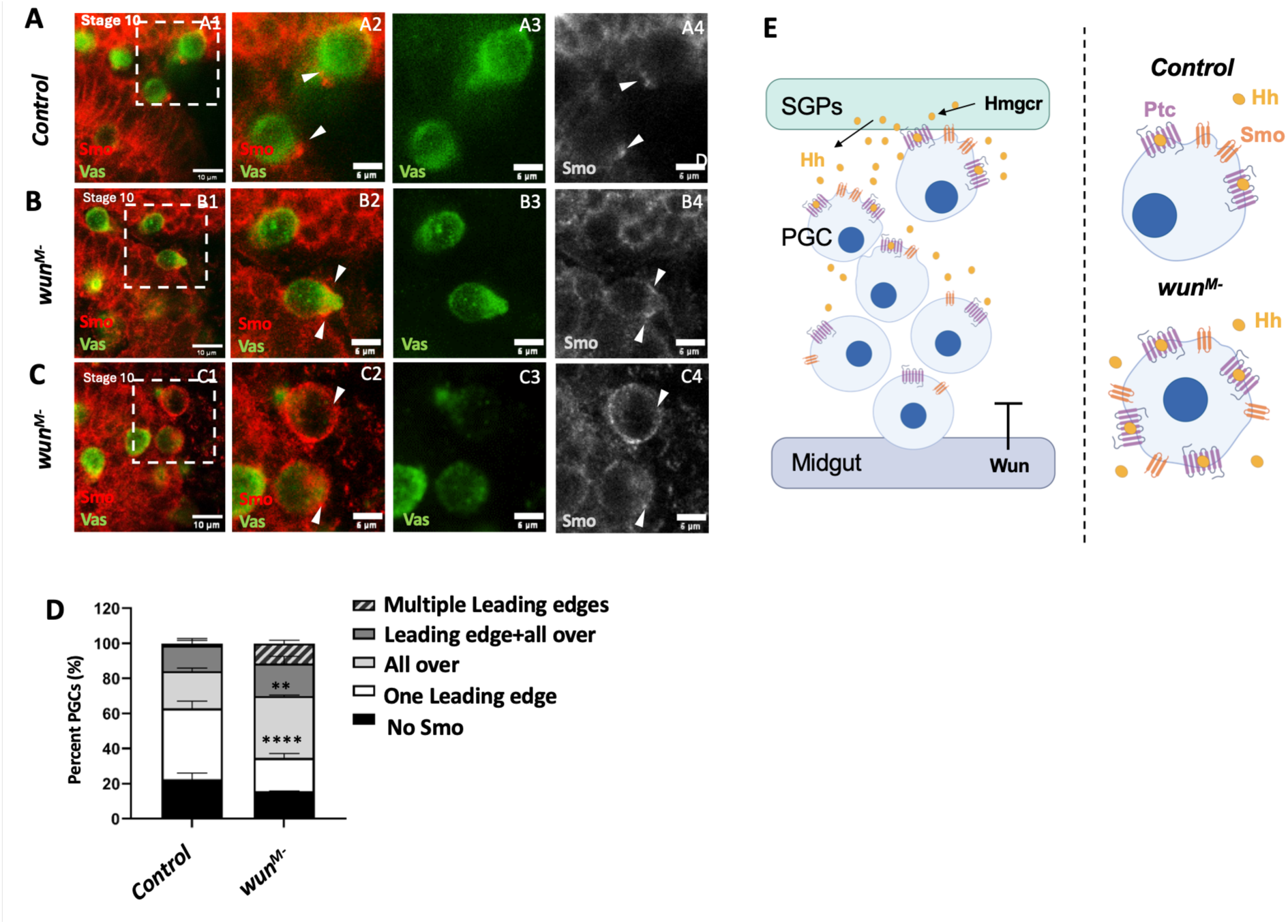
Embryos compromised for *wun* display aberrant Smo localization. **(A-E)** PGC migration through the gut in stage 10 embryos of the indicated genotype. In control *lexA^M-^*embryos (A) Smo is localized at the leading edge (A1, magnified inset A2-A4). In contrast, in *wun^M-^* PGCs, multiple foci of Smo localisation are observed (B1, magnified inset B2-B4) or are present all over the PGC (C1, magnified inset C2-C4). All panels were equally edited with minimum intensity value at 10 and maximum intensify value at 100 using Fiji Image J. **(D)** Comparison of percentages of embryos of indicated genotypes having Smo localization as per mentioned classes were quantified (n=40-70 across replicates) using 2-way ANOVA and Tukey’s multiple comparison test *** p<0.001** p<0.01. **(E)** Germ cell repulsion mediated by Wunen(s) is mediated by their ability to inhibit Hh signaling in the somatic tissues such as gut, ectoderm, nervous system. This influences the path of the migrating PGCs and guides them towards their destination i.e. SGPs, enriched in Hmgcr, which potentiates Hh activity. Additionally, germ cell-germ cell repulsion is also mediated by Wun and germ cell autonomous ability of Wun(s) to inhibit Hh signaling may be critical in this regard and upon compromising *wun* activity response to Hh signaling gets activated precociously.

### PGC migration defects in wun^M-^ embryos mimic those in ptc^M-Z+^ embryo

In earlier studies we found that PGCs in stage 11 embryos generated by mating *ptc^-^* germline clone females to wild type males tended to get stuck in the gut and/or clump together (Deshpande et al., 2001). Since loss of *ptc* would result in the constitutive activation of the *hh* signaling pathway, one might imagine that the excessive multi-directional *hh* signaling evident in *wun^M-^* embryos could result in the PGC phenotypes observed in the absence of *ptc*. To test this idea directly, we maternally depleted *ptc* and the *wun(s)* using the same RNAi strategy described above (*nos-Gal4/UAS-ptcRNAi*, *nos-Gal4/UAS-wunRNAi* and *nos-Gal4/UAS-wun2RNAi* females mated to wild type males) Suppl. Fig. 2 and Fig. 5. As controls, we generated embryos from mothers expressing *lexARNAi* (*nos-gal4*/*UAS-lexARNAi*).

**Figure 5.**
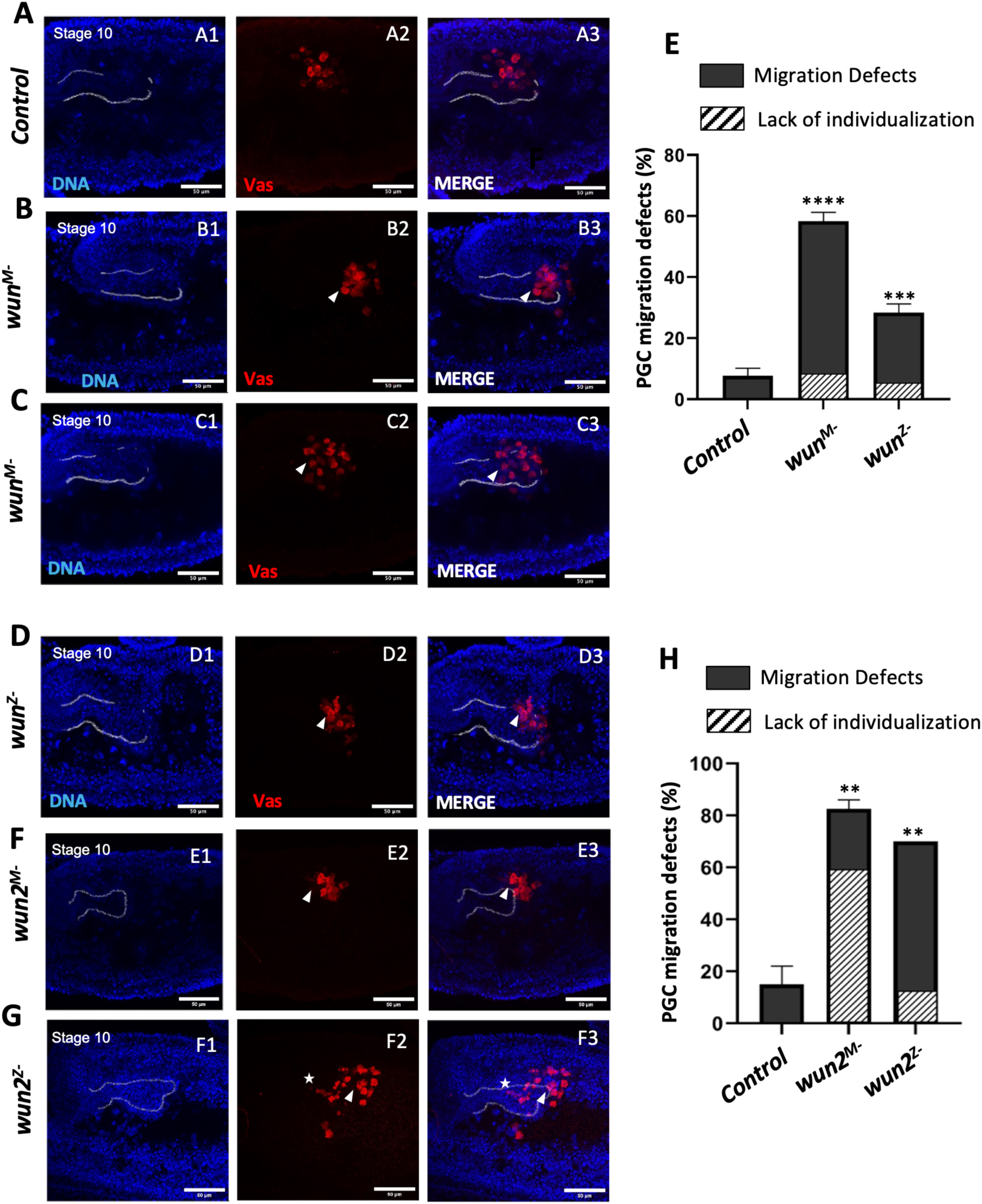
*Loss of wun* leads to PGC migration defects that are qualitatively similar to loss of *ptc*. All embryos shown are at Stage 10 **(A-D)** PGCs in control *lexA^M-^*embryo (A) exit the gut through its side and enter the mesoderm. In contrast, PGCs in *wun^M-^* embryos either clump precociously (B) or are delayed while migrating through the gut (C). PGCs of *wun^Z-^* (D) show a similar phenotype, although, as expected, the penetrance is lower. **(E)** PGC migration defects on depleting *wun* are quantified (n=15, 3 replicates) and significance determined using Ordinary one-way ANOVA and Tukey’s multiple comparison test (n=15, 3 replicates), ****p<0.0001, ***p<0.001. **(F-G)** *wun2^M-^*leads to a highly penetrant precocious clumping phenotype in stage 10 (E)*. wun2^Z-^* embryos also show similar phenotype (F) although not as penetrant. Arrowheads mark mis-migrated PGCs. Additionally, dying PGCs, marked by a star, are observed **(H)**. PGC migration defects on depleting *wun2* are quantified (n=10, 2 replicates) and significance determined using Ordinary one-way ANOVA and Tukey’s multiple comparison test (n=10, 2 replicates) **p<0.01. Hatches represent the percentage of embryos that have PGCs that have not individualized by stage10. Embryos were stained with Hoechst to mark DNA (A1-D1 and E1-F1) and anti-Vasa (A2-D2 and E1-F1). (A3-D3 and E3-F3) display merged images.

As shown in Suppl Fig. 3 we find that individualization of the PGCs in nearly half of the *ptc^M-^* embryos (45%) is hindered, resulting in an inefficient exit across the midgut primordium. Consequently, PGCs lag behind in the gut forming small clumps. In a small but significant proportion of the *ptc^M-^* embryos PGCs form a large clump inside the gut (Suppl. Fig. 2B). Similar phenotypes are observed in stage 10-13 embryos derived from *nosGal4*/ *UAS-wun RNAi* females. In >55% of *wun^M-^* embryos the individualization of PGCs was hindered, resulting in a delay in exiting the gut (Fig. 5C-G). In about 15% of embryos, the PGCs clump together within the midgut (Fig. 5B, G). Maternal depletion of *wun2* results in similar PGCs phenotypes to those observed in *wun^M-^* embryos; however, they are substantially more penetrant. Over 60% *wun2^M-^* embryos have large clumps of PGCs either within the gut or along the dorsal surface of the gut (Fig. 5F, H). In another ∼15% of the *wun2^M-^* embryos, several PGCs are delayed in exiting the gut.

### wun or wun2 are required in PGCs during the initial phase of migration

The results in the previous section indicate that the *wun(s)* play a key role in directing PGC migration at a very early step when the cells first begin to exit the midgut. Since the maternal *RNAi* knockdowns results in a substantial upregulation in the transmission of the Hh ligand throughout the embryonic soma, this early PGC defect could simply be a consequence of excessive somatic *hh* signaling. On the other hand, maternal knockdown of the *wun(s)* will also reduce the activity of the *wuns* in the PGCs themselves. In this case, the early phenotypes could also reflect a PGC-autonomous requirement of the *wun(s)*. To test this possibility, we used *nos-Gal4* to drive *UAS-wunRNAi* (Fig. 5D) or *UAS-wun2RNAi* (Fig. 5G) in migrating PGCs and stained F1 embryos with anti-Vasa. In both cases we observe phenotypes similar to those in the maternal knockdowns, though they are not as severe. For the *wun* knockdown, PGC clumping was observed in about 5% of the embryos, while lagging PGCs were observed in about 20% of the embryos (Fig. 5E). The knockdown of *wun2* was more effective: Premature clustering of the PGCs was observed in nearly 10% of the embryos, while 50% of the embryos had lagging PGCs (Fig. 5H). These findings confirm the idea that the *wun(s)* are required in PGCs for proper migration and suggest that this function is likely related to the proper functioning of the *hh* signaling pathway.

### Sac1, a known lipid phosphatase, downregulates Hh signaling in the embryonic contexts including in the PGCs

As mentioned above, the *wun(s)* are not the only phospholipid phosphatase that function to downregulate *hh* signaling. The lipid phosphatase, Sac1, was found to negatively regulate the *hh* pathway in wing and eye discs (Del Bel et al., 2018; Yavari et al., 2010). As Sac1 is also expressed in the embryonic gut primordium and the ectoderm we decided to test if maternal loss of *sac1* results in embryonic phenotypes similar to those observed for the *wuns*. We first examined the expression En in the ectoderm. Suppl Fig. 4 shows that as in the case of knockdown of the *wun(s)*, both the width of the En stripes and the nuclear signal intensity are enhanced in *sac1^M-^* embryos.

Next, we tested whether the PGCs from *sac1^M-^* embryos display PGC migration defects. We find that the PGCs in *sac1^M-^* embryos also exhibit migration phenotypes, though they are less severe than those in *wun^M-^* or *wun2^M-^*. PGCs lagging in the midgut are observed in stage 10 *sac1^M-^*embryos, and small clumps of PGCs are seen in the midgut of some stage 11 *sac1^M-^*embryos (Fig. 6; compare A, B with D, E respectively). In stage 13 embryos, there are scattered PGCs, and the number of PGCs coalescing into the embryonic gonad is reduced (Fig. 6C versus 6F). The same phenotypes are observed when *nos-Gal4* is used to drive paternally derived *UAS-sac1RNAi* in the migrating PGCs (Fig. 6; G, H vs I, J respectively). Interestingly, zygotic knockdown of the reciprocally active Kinase, *stt4*, also leads to PGC migration defects (Yavari et al., 2010); (This manuscript, Fig. 6 K-M; See Suppl. Fig. 4 for quantification)). Furthermore, supporting a genetic interaction between *wun* and *sac1*, a higher proportion of embryos displayed precocious clustering of PGCs (Fig. 6 P) and Smo mislocalization at germ band extension stage (Suppl. Fig. 5).

**Figure 6.**
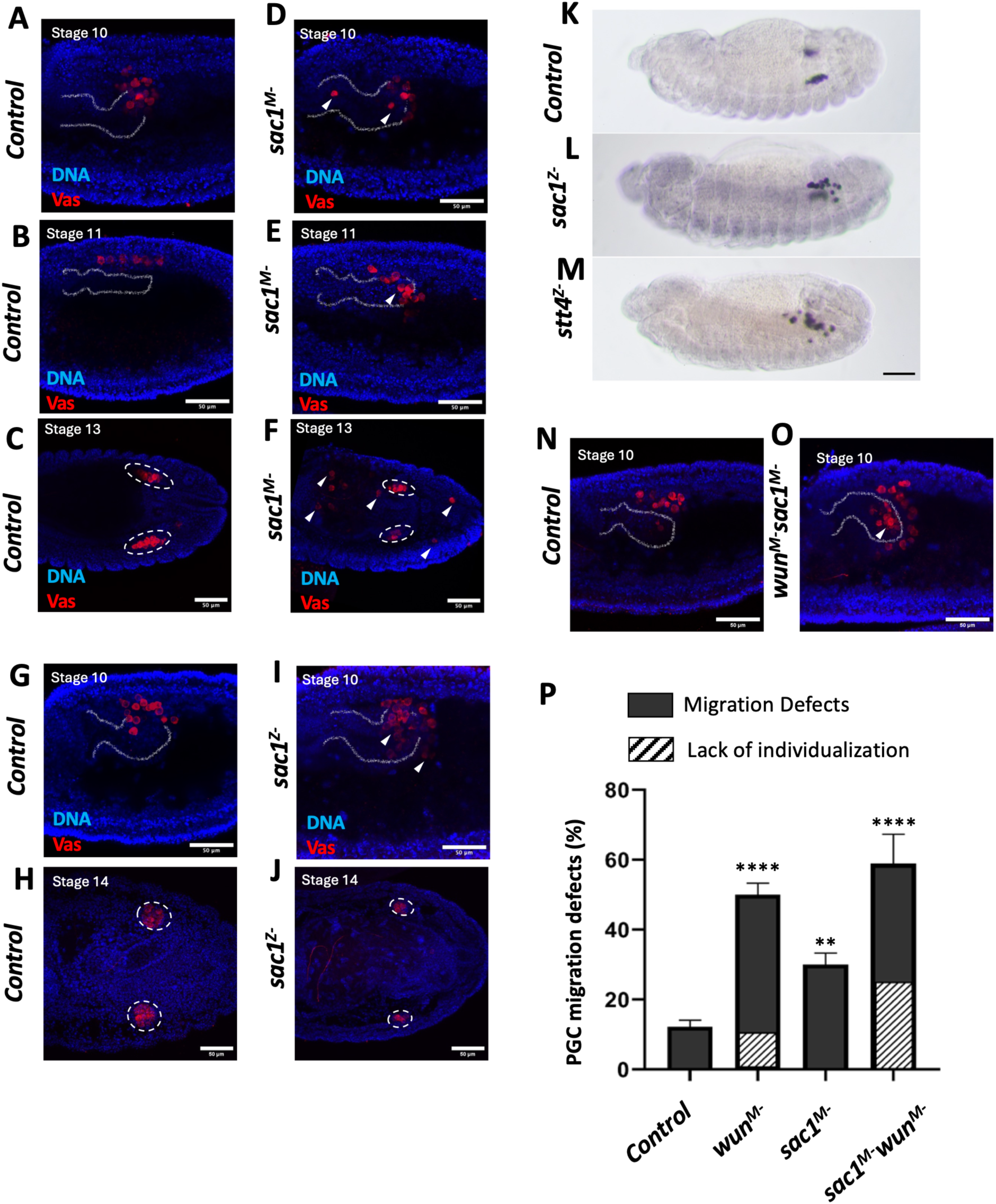
Sac1 influences PGC migration. **(A-J)** Embryos of indicated stages and specified genotypes stained with anti-Vas and Hoechst to mark DNA. (A-F) Embryos that are maternally deprived of indicated genes. Compared to control (A-C) *lexA^M-^*, the PGCs of *sac1^M-^*embryos have a variety of migration defects such as delay in exiting the gut (D), clumping (E) and inability to migrate towards the SGPs (F). **(G-J)** Embryos derived from *nos-gal4* mothers crossed to *UAS-RNAi* of indicated genes. Compared to control (G-H), PGCs from *sac1^Z-^*embryos also show migration defects including delay in exiting the midgut (I) and have reduced number of PGCs in the coalesced embryonic gonads as compared to control (J). Arrowheads mark mis-migrated PGCs. In the relevant panels outline of the gut (stage 10) or the embryonic gonad (stage 14) are marked with dotted lines and circles respectively. **(K-M)** Stage 13 embryos derived from *nos-gal4-VP16* females crossed to *UAS-RNAi* males of indicated genotypes resulting in embryos that are zygotically depleted for the individual gene expression in a PGC specific manner. *sac1* and *sst4* regulate Hh signaling in a qualitatively opposite manner. Compared to the control *egfp^Z-^* embryo (K), PGCs in *sac1^Z-^* embryos (L) are slow to reach the embryonic gonad. While PGC in *stts4^z-^* embryos scatter (M). (N-O) Stage 10 embryos which are maternally deprived of indicated genes. Compared to control (N), simultaneous maternal depletion of *sac1* and *wun* result in worsening of PGC migration defects and leads to more precocious clumping of PGCs (O). (P) Quantification of comparison of PGC migration defects (n=15, 3 replicates) and significance calculated using Ordinary one-way ANOVA and Tukey’s multiple comparison test ****p<0.0001, ***p<0.01. Diagonal lines represent the percentage of embryos that have PGCs that have not individualised by stage10. The gut outline is marked. The gonads at stage 13-14 are marked within dashed circles.

### Endogenous Hh-GFP specific puncta in the mesoderm and the adjacent PGCs are enhanced in *sac1^M-^*embryos

The analysis of *sac1* function in the eye and wing discs suggested that it functions in receiving cells to inhibit the membrane association of Smo (Griffiths et al., 2020; Yavari et al., 2010). When *sac1* activity was compromised in receiving cells, Smo association with the membrane was substantially enhanced. Moreover, in this developmental context the phenotypes induced by the loss of *sac1* activity were found to be independent of the Hh ligand. Consistent with a similar cell-autonomous role, we found that knockdown of *sac1* activity in PGCs induced migration defects. However, since we also observed PGC migration phenotypes when *sac1* activity is maternally depleted, it seemed possible that *sac1* might also have a cell non-autonomous function in the *hh* signaling pathway analogous to *wun(s)*. To explore this possibility, we mated *Mat Gal4*/*UAS-sac1-RNAi* females with *hh-GFP* males. The experiments in Fig. 7 show that loss of *sac1* results in a substantial upregulation in the transmission of the Hh ligand. In panels A-C (Fig. 7) we examined the accumulation of Hh-GFP puncta in the mesoderm. As indicated by the quantitation in panel C, the number of Hh-puncta is increased ∼2.5 fold. Accompanying this increase in mesodermal Hh-GFP puncta, we find that there is a significant increase in the number of Hh-GFP puncta per PGC in *sac1^M^*^-^. (Fig. 7F). About 55% control PGCs had 1-3 Hh-GFP puncta as opposed to ∼20% of the *sac1^M^*^-^ PGCs. ∼ 80% of *sac1^M^*^-^ PGC had 4 or more Hh-GFP puncta compared to about 53% in control. More importantly, none of the PGCS in control embryos had 10+ Hh-GFP puncta. Altogether, our data shows that maternally compromising *sac1* activity results in excess accumulation of Hh-GFP in the mesoderm and PGCs.

**Figure 7.**
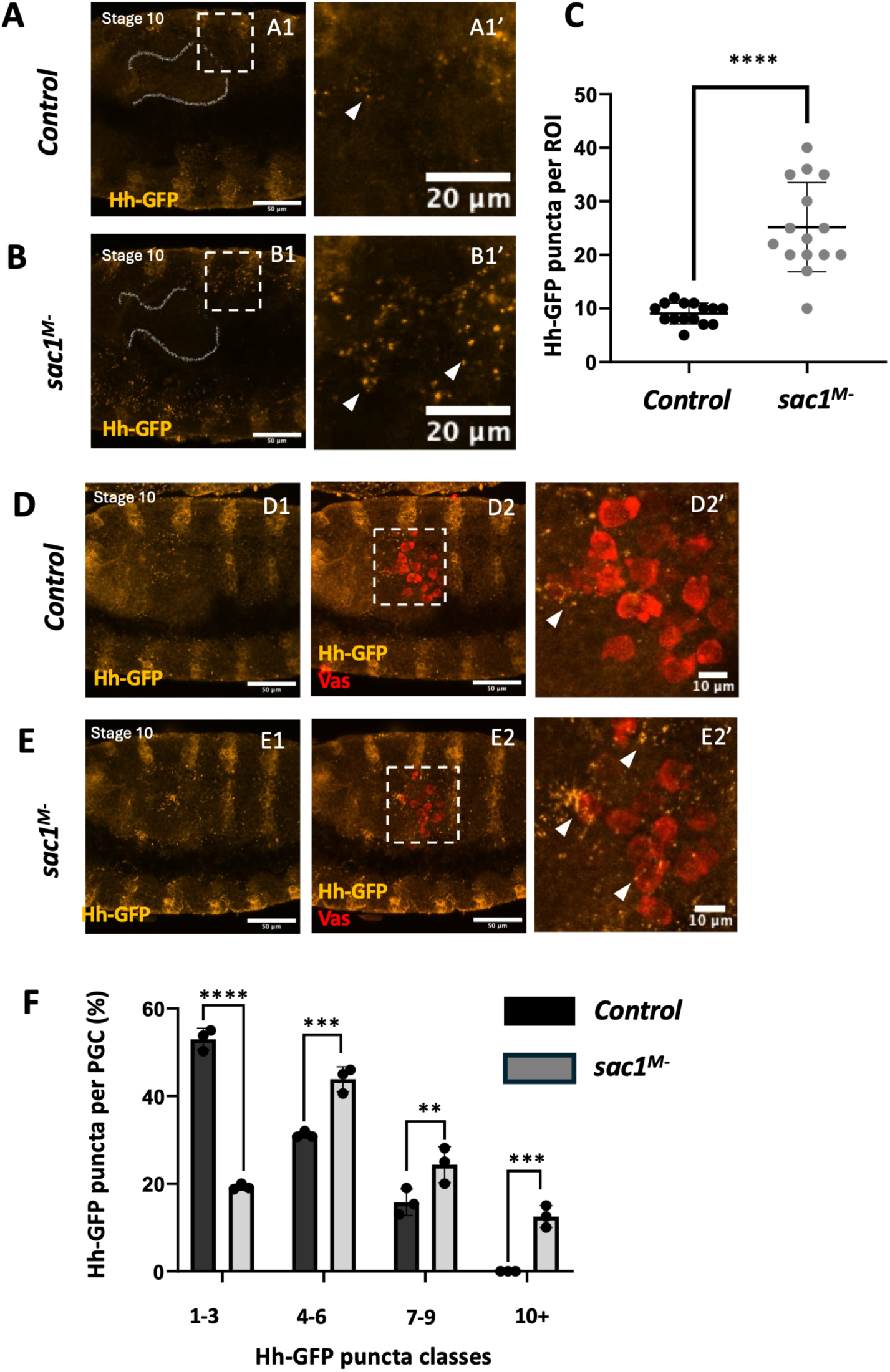
Endogenous Hh-GFP specific puncta in the mesoderm and the adjacent PGCs are enhanced in sac1^M-^ embryos. **(A-B)** Either *UAS-lexA RNAi* (control) or *UAS-sac1 RNAi* males were mated with *MatGal4* females. Virgin females carrying both the transgenes were subsequently mated with males expressing endogenous *hh-GFP*. Embryos resulting from this cross are maternally depleted of indicated genes, express endogenous *Hh-GFP* and are referred to as *lexA^M-^* or *sac1^M-^*. Compared to control (A) *sac1^M-^* embryos show higher number of *Hh-GFP* puncta (yellow) in the mesoderm(B). **(C)**The increase in *Hh-GFP* is quantified using an unpaired T-test. Individual *Hh-GFP* puncta were counted in 15 ROIs above gut, marked, per genotype. ****p<0.0001. (A1-B1) Embryos stained with anti-GFP (Yellow). Magnified insets are shown in (A1’-B1’). **(D-E)** Stage 10 embryos maternally depleted of the indicated gene, expressing endogenous Hh-GFP. Compared to control *lexA^M-^* (D), *sac1^M-^* embryos show higher number of *Hh-GFP* puncta in their PGCs. Hh-GFP stained with anti-GFP antibody (D1-E1, yellow) along with PGCS stained with anti-Vas (D2-E2, red). **(F)** Total number of puncta were counted in 12 PGCS per genotype across three replicates. The percentages of PGCs with 1-3, 4-6, 7-9 and 10+ *Hh-GFP* puncta are indicated in the graph and significance calculated using Two-way ANOVA and Dunnett’s multiple comparison test. ****p<0.0001, **p<0.01. (D2’-E2’) are magnified insets. Arrowheads mark Hh-GFP puncta.

### Comparative unbiased lipidomics of wun^M-^ and sac1^M-^ embryonic extracts

While depletion of the *wun(s)* and *sac1* give a similar range of *hh* signaling pathway phenotypes, the substrates for these enzymes involved in lipid metabolism are likely different. The preferred substrate for *sac1* is phosphatidylinositol 4-phosphate (PI4P) which it dephosphorylates to phosphatidylinositol (PI) (Yavari et al., 2010).

While the *wun(s)* share homology with a mammalian family of lipid phosphate phosphatases (LPP1, LPP2 and LPP3), biochemical experiments suggest that their substrate preferences are likely different. The mammalian LPP family proteins dephosphorylate a broad range of phospholipids, including among others lysophosphatidic acid (LPA), phosphatidic acid (PA) and ceramide-1-phosphate C(1)P (Dillon et al., 1997; Jasinska et al., 1999). Unlike the mammalian LPP family members, (Burnett and Howard, 2003) found that *wun* is unable to dephosphorylate either PA or C(1)P. However, *wun* was able to dephosphorylate LPA with similar efficiency to the mammalian LPP1 and LPP3 enzymes. To identify possible targets for the *wun* and *sac1* we performed unbiased lipidomics on embryonic extracts derived from *wun^M-^* and *sac1^M-^* 4-8 hr embryos (for details, see Materials and Methods).

Because compromising *sac1* has been reported to increase PI4P levels, we initially focused on phosphoinositide levels. Notably, embryos maternally compromised for *sac1* showed increased levels of various phosphoinositide species (estimated and collectively represented as PI) compared with control samples. Evidently, PI levels are not appreciably altered in *wun^M-^* embryos. Surprisingly, however, DAG levels are considerably elevated in *wun^M-^* extracts as compared to either the control or the *sac1^M-^* samples. Membrane-bound DAG subsequently participates in a variety of physiological alterations (Falzone and MacKinnon, 2023; Nilssen et al., 2005; Zhong et al., 2022).

Since DAG is not phosphorylated, its accumulation in *wun^M-^* extracts cannot be attributed directly to the loss of *wun* activity. Instead, the elevated accumulation of DAG must be an indirect consequence of the loss of *wun* function. Elevated levels of DAG are typically generated by the activation of phospholipase C, which breaks down PIP_2_ into DAG and IP3. Typically, signaling pathways that utilise GPCRs can turn on phospholipase C by promoting the dissociation of *Gα* from the Gβγ dimer. Phospholipase C can also be activated by excessive levels of lysophosphatidic acid (LPA) which is the only known *wun* substrate (Burnett and Howard, 2003). Thus, the specific and robust increase in DAG levels upon compromising Wun activity dovetails nicely with aberrant localization of Smo, a GPCR involved in mediating Hh signal. Together these data support the notion that Wunen, likely employs Hh and the downstream components of the signaling cascade to engineer germ cell repulsion.

## Discussion

Migrating cells in different developmental contexts are exposed to a plethora of signals in their cellular environment. Only a select few of these signals has the ability to guide motility of these cells and can function as either attractive or repulsive cues. Conversely, the migrating cells are also endowed with a restricted and well-calibrated ability to respond selectively to either move away from or towards the signal. For instance, axonal migration in vertebrate embryos is guided by attractive influence of Netrins and Sonic-hedgehog whereas they are repelled by BMP ligands (Butler and Dodd, 2003; Charron et al., 2003; Tessier-Lavigne, 1994; Wu et al., 2019). PGC migration in *Drosophila* is engineered by Hmgcr and Wun dependent attractive and repulsive influences respectively. While Hh acts as an attractive ligand for migrating PGCs, no specific repulsive signal has been identified thus far. It has been proposed that components of the Hmgcr dependent enzymatic pathway harness attractive potential of SGP derived Hh signal. Data presented here document that repulsive influence mediated by *wun* also targets Hh, however, in a qualitatively opposite, antithetical manner. In the following, we will discuss how a competitive reciprocal balance between *hmgcr* and *wun* modulates Hh activity to carve the precise path of migrating PGCs.

The functioning of the *hh* signaling pathway in guiding the migration of the PGCs towards the SGPs is now reasonably well understood. Hh expressed by the SGPs is received by the migrating PGCs and its reception is transduced into their directed motility towards the source of the Hh. The response to the reception of the Hh ligand is highly polarized. Consequently, Smo is translocated from the cytoplasm to the membrane where it accumulates in a single sharp focus at the leading edge of the migrating PGCs. Smo subsequently recruits the GPCR Tre1 to the membrane and Tre1 activates F-actin assembly at the leading edge through the production of the signaling molecule PI(4,5)P_2_ (Kim et al., 2021).

To ensure that Hh ligands produced by the SGPs are preferentially received by the PGCs is especially critical to the deployment of *hh* as a guidance molecule in this context. Previous studies identified one such mechanism, namely the *potentiation of Hh transmission*. Though *hh* is widely expressed in ectodermal and mesodermal tissues throughout much of the embryo, its ability to attract migrating PGCs is primarily limited to the Hh produced by the SGPs (and their progenitors). One critical factor that distinguishes Hh produced by the SGPs from Hh expressed elsewhere in the embryo is *hmgcr*. *Hmgcr* and the downstream isoprenoid biosynthetic pathway function to greatly enhance the long-distance transmission of the Hh ligand. When *hmgcr* is expressed in other cell types or tissues, the endogenous Hh ligand expressed in these cells (or tissues) is able to compete with the Hh ligand produced by the SGP and induce the mismigration of the PGCs. Thus, key to the potentiation activity of *hmgcr* is the fact its expression becomes progressively restricted to the SGPs during PGC migration. In addition to *hmgcr* several other genes important for the transmission are expressed in the mesoderm or posterior mesoderm (Deshpande et al., 2016; Deshpande et al., 2013; Ricardo and Lehmann, 2009). However, the potentiation of Hh transmission from the SGPs (and their mesodermal progenitors) by *hmgcr* is not in itself sufficient. We show here that the long-distance transmission of *hh* expressed in other cell types and tissues in the embryo must also be specifically suppressed in order to ensure that the migrating PGCs make their way to the SGPs. The suppression of *hh* signaling is mediated by the two Wun(s), Wun and Wun2. It was initially discovered that Wun functioned to repel migrating PGCs (Zhang et al., 1996; Zhang et al., 1997). Subsequent studies identified the neighboring gene *wun2* (Hanyu-Nakamura et al., 2004; Starz-Gaiano et al., 2001) and showed that it also functions to repel migrating PGCs. The two *wun(s)* encode lipid phosphate phosphatases and exhibit similar patterns of expression. Both gene products are maternally deposited and are present in the soma as well as the PGCs. In addition, both genes are zygotically expressed in the epidermis, gut primordium, and nervous system, tissues that migrating PGCs usually avoid.

While the repulsive activity of the wun(s) has been well documented, the mechanism of their action was not known. Using both the ‘loss’ and ‘gain-of-function’ experiments here we demonstrate that Wun(s) “repel” migrating PGCs because they are able to inhibit Hh signaling. There are several interesting possibilities where Wun(s) may exert their effect. For instance, they could influence either the synthesis or stability of Hh ligand. Alternatively, Wun(s) could suppress the transmission of the Hh ligand—and this suppression of Hh transmission from other cell types and tissues enhances and refines the ability of Hh produced by the SGPs to attract the migrating PGCs. Perhaps the epistasis experiment presented in the manuscript constitutes the most direct demonstration of this mechanism of action involving antithetical activities of Hmgcr and Wun on Hh transmission. When *hmgcr* is ectopically expressed in the CNS, it induces PGC mismigration because it substantially enhances the transmission of the Hh ligand that is normally produced in the CNS. For this reason, the migration defects induced by ectopic *hmgcr* can be suppressed by co-expressing *hh-RNAi* with *hmgcr.* A similar suppression is observed for *wun*—in this case when *wun* is co-expressed with *hmgcr* using the same CNS driver. Future analysis will focus on determining the mechanisms employed by these lipid metabolism enzymes to inhibit Hh signaling from the SGPs to the PGCs.

Consistent with an ability to antagonize the *hh* signaling pathway, we find that when either *wun* or *wun2* is maternally depleted the *hh* signaling pathway is substantially upregulated. This enhanced activity is evident in the pattern of En expression in the embryonic ectoderm during mid-embryogenesis: there is an increase in the level of En expression and in the number of cells expressing En. Visualization of Hh produced by a BAC:*hh-GFP* rescue construct indicates that the transmission of Hh is upregulated when maternal *wun* is depleted. In a wild-type background Hh-GFP puncta decorate a subset of the migrating PGCs. In *wun^M-^* embryos, the number of Hh-GFP puncta/PGCs increases. Excess Hh-GFP puncta are also seen in the mesoderm, indicating that there is a general enhancement in Hh accumulation. The reception of the Hh ligand in migrating PGCs is also substantially altered. Instead of there being one primary membrane associated Smo focus, nearly half of the PGCs in *wun^M-^* embryos have high levels of Smo all around their membranes or multiple Smo foci.

The enhanced transmission of the Hh ligand when *wun* is maternally depleted together with the *hmgcr/wun* epistasis experiments suggest that the Wun(s) function to mitigate the *hh* signaling pathway in *hh* expressing cells. On the other hand, the Wun(s) are known to be required not only in the soma but also in the PGCs themselves. In this case, the mislocalization of Smo in PGCs of *wun^M-^* embryos could be due at least in part to an inappropriate cell autonomous activation of the *hh* signaling pathway in the receiving cells. As cytonemes extend from both Hh producing and Hh receiving cells in the imaginal discs, it is also conceivable that the loss of maternal *wun* in PGCs contributes to the increase in their labeling by Hh-GFP puncta. Two lines of evidence provide additional support for the idea that the *hh* pathway is upregulated in both the sending and receiving cells. The first would be the finding that *ptc*-like PGC migration defects are observed in both maternal *wun/wun2* RNAi knockdowns and in *wun/wun2* RNAi knockdowns in migrating PGCs. The second is the finding that a loss of another lipid phosphatase, *sac1*, also induces a similar spectrum of PGC migration defects. Consistently, as in the case of *wun*, embryos compromised for *sac1* display PGC migration defects between stages 10-13 including retention of PGCs in the gut and reduction of PGC count in the coalesced gonads in a subset of embryos. Furthermore, as observed in *wun^M-^*embryos, *sac1^M-^* embryos also display an increased Hh association with migrating PGCs and an overall increase in the level of Hh in the surrounding mesoderm.

Analogous phenotypic consequences of the embryonic knockdowns of *wunen(s)* and *sac1* and their respective influence on Hh signaling on the embryonic PGCs are interesting in their own right. These observations coaxed us to revisit the reportedly crucial involvement of lipid metabolism and function vis-a-vis Hh signaling, PGC migration and their functional relationship. Wunen(s) have been characterized as lipid phosphate phosphatases (LPPs) based on their homology to the vertebrate LPPs. Substantial genetic characterization including rescue experiments has further supported the notion that LPP3 (human) and Wunen(s) likely share targets. By comparison, Sac1 has a relatively narrower and specific target range. Especially in the context of Hh signaling, it balances the Stt4 kinase activity to maintain the levels of PI4, which in turn, regulates membrane localization of Smo to modulate signaling downstream of Hh. Consequently, the phenotypic similarities between *wun* and *sac1,* in the context of PGC migration are intriguing.

Hence, we decided to assess whether they shared targets that may explain the similar phenotypes. The comparison of unbiased Lipidomics confirmed a few expectations but also presented some unanticipated surprises. Foreseeably levels of different phosphoinositide species were substantially elevated upon the embryonic loss of *sac1*. Surprisingly however, a similar loss of *wun* did not alter PI or PA levels rendering them unlikely to be biologically relevant entities. Our current evidence supports the notion that Wunen(s) may function more broadly as a lipid metabolic or lipid-modifying enzyme(s), rather than lipid phosphatases per se. In this regard it is noteworthy that a specific and significant increase in DAG levels was observed upon loss of *wun*. The possible mechanistic underpinnings of this increase and its relationship with the phenotypic consequences are tantalizing as DAG accumulation is one of the indicators of activation of GPCR signalling (Falzone and MacKinnon, 2023; Nilssen et al., 2005; Zhong et al., 2022). Of note, Smo and Tre1, the two receptors demonstrated to act downstream of Hh signaling during PGC migration, are members of the GPCR family. Precocious clumping observed upon loss of Wun(s) could thus be an outcome of premature activation of one (or both) of these receptors.

*Drosophila* PGCs migrate in a calibrated manner across a complex cellular landscape. To arrive at their destination, germ cells must be able to respond to the ‘right’ signal and ignore the ‘wrong’ ones. Moreover, PGCs must distinguish between the right signal emanating from the wrong source. Wunen(s) contribute to this ability as they constitute the other side of the coin. They hinder Hh’s ability to function as an attractive force if it emanates from a ‘wrong source’.

The antithetical abilities of Hmgcr and Wun thus allow fine-tuning of Hh signalling by potentiating the SGP-derived ‘right’ signal and squelching the one emanating from the ‘wrong’ sources like ectoderm, nervous system, etc. Moreover, this is unlikely to be an isolated instance and thus harbors tremendous potential as various disease conditions including metastasis involve long distance migration and inappropriate adhesion and lodging. Future experimental strategies will help elucidate the mechanistic underpinnings of these events in the context of PGC migration and may also pave the way for understanding its importance in other contexts where directed cell migration is a critical determinant of either normal or aberrant development.

## Materials & Methods

### Fly husbandry and stocks

Flies were raised on standard cornmeal agar at 25 °C unless stated otherwise. The *Drosophila melanogaster* lines, with IDs, used are: *CS* (BDSC_64349), *UAS-lexA RNAi* (BDSC_67947), *UAS-eGFP* (BDSC_41552)*, UAS-ptc RNAi* (BDSC_55686), *UAS-smo RNAi V20* (BDSC_53348), *UAS-smo RNAi V22* (BDSC_43134), *UAS-wun RNAi* (BDSC_32429 and 32430) and *UAS-wun2 RNAi* (BDSC_32423), *nos-gal4 VP16* (BDSC_4937)*, UAS-sac1 RNAi* (BDSC_56013), *UAS-sac1 RNAi (* VDRC_44376 (L1) and VDRC_37216 (L2)), *UAS-mcherry* RNAi (BDSC_35785), *maternal-〈-tubulin-gal4-VP16* (*Mat-gal4 on both II and III*, Gift from Daniel St. Johnston), *ptc-gal4 UAS-lacz* (Gift from Matthew P Scott), *Hh:GFP Bac* (Gift from Thomas B. Kornberg), *UAS-wun-GFP* (Gift from Ken Howard).

For maternal knockdown experiments, we crossed *UAS-X RNAi* males, where ‘X’ is the gene to be knocked down, to maternal gal4 driver females (either *mat-gal4 VP16 or nos-gal4 VP16*). F2 embryos were collected for study, and since these are maternally compromised, their genotype is denoted as *X^M-^.* For zygotic knockdowns, denoted as X^Z-^, *UAS-X RNAi* males were crossed to maternal gal4 drivers (*mat-gal4* or *nos-gal4*) females, and F1 embryos were used for study. *nos-gal4 VP16*, unlike *mat-gal4 V16* is enriched specifically in PGCs. For zygotic overexpression of *wun*, we used F1 embryos (denoted as *wun^OE^*) derived from *mat-gal4* mothers crossed to males carrying the *UAS-wun-GFP* transgene.

### Embryo Fixation

Embryos were collected, typically aged 3-8 hours (with a focus on stages 10-11) or 0-12 hours (for overnight collections of later stages), in a sieve and dechorionated in 4% sodium hypochlorite for 90 seconds. After thorough washing with distilled water, the embryos were fixed in a 1:1 heptane:4% Paraformaldehyde (PFA) solution for 20 minutes, followed by removal of the PFA and a fresh heptane wash for 20 minutes. Furthermore, chilled methanol was added, and the embryos were devitellinized by vigorous shaking in a 1:1 heptane: methanol mixture. Devitellinized embryos settle at the bottom and are stored in 100% methanol at –20 °C until immunostaining.

### Immunostaining and Imaging

For immunostaining, embryos were rehydrated by four washes with 0.3% PBS-Triton X-100 (PBST), followed by blocking in 2% bovine serum albumin (BSA) in 0.3% PBS-T (blocking solution) for 1 hour at room temperature. Embryos were incubated overnight at 4°C with primary antibodies diluted in fresh blocking solution at the appropriate antibody-specific dilutions. The following primary antibodies in specified dilutions were used: Rabbit anti-Vasa (1:1000, Gift from Trudi Schüpbach), Mouse anti-En (1:50, DSHB), Mouse anti-Wg (1:20, DSHB), Chicken anti-GFP (1:500, Invitrogen), Mouse anti-Smo (1:50, DSHB), Mouse (1:20), and Chicken anti-®gal (1:1000, Abcam). Following four washes with 0.3% PBS-T, the embryos were incubated in the appropriate secondary antibodies for 1 hour at room temperature. The following secondary antibodies were used in a 1:1000 dilution: goat anti-mouse AlexaFluor 488, goat anti-chicken AlexaFluor 568, goat anti-rabbit AlexaFluor 647 (1:1000, Invitrogen). Embryos were washed four times with 0.3% PBS-T, and Hoechst (1:1000 from a stock of 10 mg/ml) was added to the penultimate wash. All washes mentioned in the protocol are 10-15 mins each. Embryos were mounted in 20mM Tris pH=8, 0.5% n-propyl gallate, 90% glycerol and observed under a Nikon AX confocal microscope or an Evident inverted microscope (FV4000) using 40X and 20X objectives, respectively. Image processing was performed using the ImageJ program.

### Lipidomics

*Mat-Gal4* females were crossed with each of *UAS-mCherry RNAi*, *UAS-wun RNAi, and UAS-Sac1 RNAi* (two lines, *L1 &* L2). F2 embryos were collected for 0-4 hours and then aged at 25 °C for 4 hours. Embryos were washed in 1x phosphate buffered saline (PBS) and snap frozen until further processing. For lipid extraction, the embryos were thawed on ice, suspended in 550 μL of cold, sterile 1x PBS, pH 7.4 and homogenized using a tissue homogenizer (Bullet Blender24, Next Advance) with one scoop of glass beads (0.5-mm diameter; Next Advance) at a speed setting of 10 for 5 min at 4 °C. The protein concentration of each lysate was estimated using the BCA protein assay kit (Pierce) according to the manufacturer’s protocol. 500 μL of the lysates were used for lipid extraction using established methods as described earlier (Kumar et al., 2024). Briefly, for each sample, 500 μL lysate was transferred to a glass vial and vigorously mixed with 1.5 mL of 2:1 chloroform (CHCl_3_): methanol (MeOH) containing an internal standard mix [*1 nmol each of pentadecanoic acid (Sigma-Aldrich, Catalog No. P6125, for negative ion mode analysis) and C17:0/20:4 phosphatidylcho line (Avanti Polar Lipids, Catalog No. LM-1002, for positive ion mode analysis)*]. The resulting mixture was centrifuged at 1500g for 15 min to separate the aqueous (top) and organic (bottom) layers, and the organic layer (bottom) was gently pipetted into a new glass vial. To enrich the extraction of phospholipids, the aqueous layer was acidified using 20 μL formic acid (Merck, Catalog No. 5.33002) and further vigorously mixed with 1 mL of CHCl_3._ The resulting mixture was again separated by centrifugation at 1500g for 15 min, and the organic layer (bottom) was separated and pooled with the organic extract from the previous extraction. The pooled organic phase was dried under a stream of nitrogen gas, resolubilized with 500 μL of CHCl_3_, and transferred to another glass vial to remove any trace aqueous contamination. The organic layer was dried again under a stream of nitrogen gas and resolubilized in 200 μL of 2:1 CHCl_3_/MeOH, and 10 μL of this was injected into an Agilent 6545 Quadrupole Time-of Flight (QTOF) LC-MS instrument for untargeted lipidomics measurements in each ion mode (positive and negative). A standard untargeted lipidomics run in positive or negative ion mode lasted 60 min, and the LC columns, solvents, and solvent gradients were identical to those previously reported (Kumar et al., 2024). All LC-MS/MS experiments were performed using an electrospray ionization (ESI) source with the following parameters: drying and sheath gas temperature = 320 °C; drying and sheath gas flow rate = 10 L/min; fragmentor voltage = 150 V; capillary voltage = 4 kV; nebulizer pressure = 45 psi; and nozzle voltage = 1 kV. For the analysis of different lipids from the embryos, a Personal Compound Database Library (PCDL) curated for lipids was generated, and peaks corresponding to lipids were verified based on relative retention times, mass error threshold (20 ppm or less at MS1), and MS/MS fragments obtained. All lipid species were quantified by measuring the area under the curve relative to the respective internal standard and then normalizing this to the protein concentration of the respective sample.

### Quantification & Statistical Analysis

The statistical analysis are detailed in figure legends. n in each graph represents the number of embryos, and N represents the number of replicates. To quantify PGC migration defects, embryos were stained with α-Vasa antibody and imaged. In embryos older than stage 13, PGCs away from the somatic gonads were counted as mis-migrating cells. For maximum-intensity projections, equivalent slices were used across genotypes to ensure the best representation.

## Contributions

GD conceptualized the project, with inputs from PS, SK and GSR. AR executed the experiments, with contributions from AER, JD, AI and GD. KK conducted the Lipidomics experiments under SK’s supervision. GD wrote the initial draft, with input from AR, AER, KK, SSK, and GSR. All authors edited and refined the manuscript to its final form. GSR, GD, PS & SSK were responsible for funding acquisition, supervision and project management.

## Acknowledgements

We thank: Bloomington *Drosophila* Stock Centre (BDSC), supported by NIH grant P40OD018537, for fly stocks; Snehal Patil and Yashwant Pawar for fly media and stock maintenance; Microscopy facility at IISER Pune, managed by Dr Santosh Podder and Vijay Vitthal; Saddam Shekh for technical assistance and maintenance of the biological mass spectrometry facility at IISER Pune.

## Supplementary Files

**Supplementary Figure 1. (A-C)** Recombinant *ptc-gal4 UAS-β-gal* females were mated with the *UAS-RNAi* males of the indicated genotype. Stage 10/11 embryos (F1) derived from the respective crosses were stained with anti-*β-gal* antibody. Compared to the *control, ptc-gal4 UAS-β-gal* (A), all embryos *ptc-gal4 UAS-β-gal/ UAS-RNAi* (n∼30) show considerably enhanced β*-gal* expression, which indicates Ptc activation (B and C). Panel B & C are biological replicates.

**Supplementary Figure 2. Maternal requirement of Hh receptors, *ptc*, and *smo* during PGC migration in early embryos.** (A-C) Stage 10 embryos maternally deprived of indicated genes were laid by the nos-gal4>UAS-RNAi of indicated genes. PGCs in control *lexA^M-^* embryo display clearly individualized PGCs that traverse across the gut through its dorsal side (A). In contrast, PGCs in *ptc^M-^* embryos either clump and fail to individualize altogether (B) or show delayed migration through the gut possibly due to incomplete individualization. Conversely, PGCs of *smo^M-^* (C) are able to individualize like the control PGCs and exit the gut but start to scatter on their way to the mesoderm. Panels (A1-C1) display Hoechst staining, Panels (A3-C3) displayed merged images. Arrowheads mark mis-migrated embryos. Gut is outlined. (D) Aberrant migration of PGCs is quantified in graph using Ordinary one-way ANOVA and Tukey’s multiple comparison test (n=12,3 replicates), ****p<0.0001. Diagonal lines represent percentage of embryos that have PGCs that have not individualized by stage10.

**Supplementary Figure 3. Embryos derived from mothers compromised for *sac1* (*sac1^M-^*) show elevated En expression in the ectoderm.** (A-B) Stage 10 Embryos derived from *nos-gal4>UAS-RNAi* were stained with anti-Engrailed (En) antibody (green) and DNA dye, Hoechst (not shown*). Compared to the negative control *lex^M-^* (A1 and A2) embryos a significantly higher percentage of *sac1^M-^* (B1 and B2) embryos display expansion in the width of the individual stripes and increased level of En protein and. ‘+’ mark is used for the cells positive for En in a stripe. Panels (A1’,-A2’ and B1’-B2’) show magnified insets. (C-D) Quantification of En stripe broadening (C) and increased En expression (D) in embryos. Significance calculated using Ordinary one-way ANOVA (n=10 N=3), ****p<0.0001.

**Supplementary Figure 4. Zygotic depletion of lipid modification enzymes required for regulating Hh signalling results in germ cell migration defects**. Bar graph showing the percentages of embryos of indicated genotypes with a given number of mis-migrating germ cells. Chi-square Goodness of Fit test was done for each genotype with the *Contro*l (n = 38), sac1^Z-^(n = 48) p = 2.01 x 10^-54^, stt4^Z-^(n = 32) p = 4.83 x 10^-**23**^

**Supplementary Figure 5. Genetic interaction between *wun* and *sac1* enhances aberrant Smo localization.** (A-C) Stage 10 embryos of the indicated genotype. In control *lexA^M-^*embryos (A) Smo is localized at the leading edge (A0, magnified inset A1-A3). In contrast, in *sac1^M-^; wun^M-^* PGCs, multiple foci of Smo localisation are observed (B0, magnified inset B1-B3) or are present all over the PGC (C0, magnified inset C1-C3). All panels were equally edited with minimum intensity value at 10 and maximum intensify value at 100 using Fiji Image J. (D) Comparison of percentages of embryos of indicated genotypes having Smo localization as per mentioned classes were quantified (n=40-70 across replicates) using 2-way ANOVA and Tukey’s multiple comparison test *** p<0.001** p<0.01.

**Supplementary Data 1. Lipidomics dataset.** Prism file containing the raw data that was used to plot the heat map shown in Figure 8. The *Mat-gal4* driver was used for these experiments (see Materials & Methods), and *UAS-mCherry* knockdown embryos were used as controls. Within the file, columns (groups) A-F represent biological replicates for control experiments, and columns G-L represent *wun^M-^*; columns M-Q & R-W represent replicates of maternal knockdowns for two independent *Sac1* lines (*sac1^M-^*).

**Figure 8.**
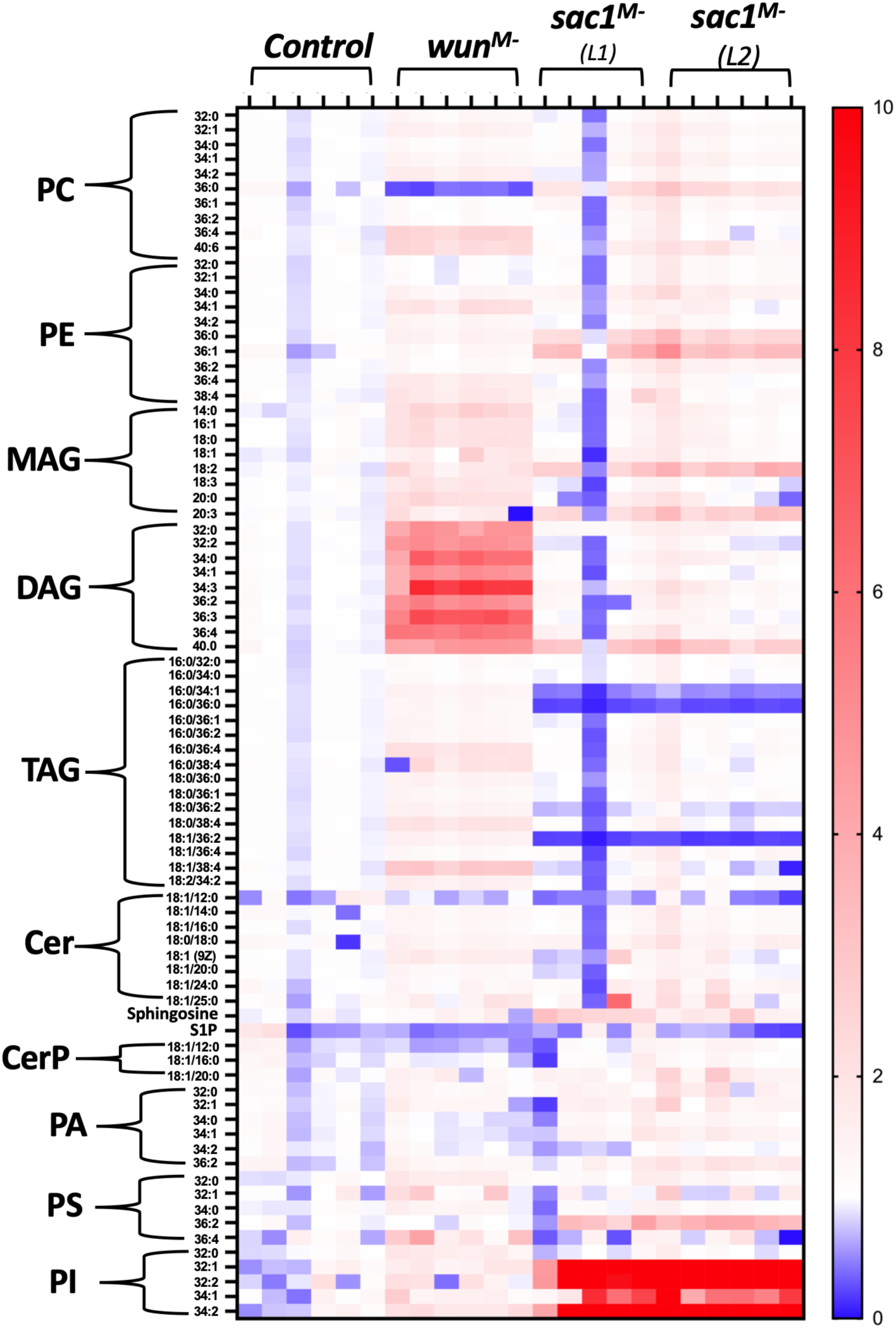
Substrate specificity of Wun and Sac1. A heat map plot showing relative lipid levels in 4-8 hour old whole embryos, post-fertilisation after maternal depletion of *wun* and *sac1*. The *Mat-gal4* driver was used for these experiments (see Materials & Methods), and *UAS-mCherry* knockdown embryos were used as controls. Lipidomic analysis shows that loss of *wun* leads to a specific increase in DAGs, whereas reducing *sac1* results in a considerable accumulation of PIs. Sac1I is known to dephosphorylate Phospho-isoforms of Phosphatidylinositols to generate PIs with a reduced number of phosphate residues. The Y-axis represents the lipid classes identified in the embryonic lysates, with individual rows corresponding to different attached fatty acids, listed by chain length and degree of (un)saturation. The X-axis represents multiple replicate experiments for the genotypes shown, with two independent *Sac1* knockdown lines used.

