## Supplementary material for "Wunen(s) help navigate Primordial Germ Cells by attenuating Hedgehog signaling": Suppl. Figures

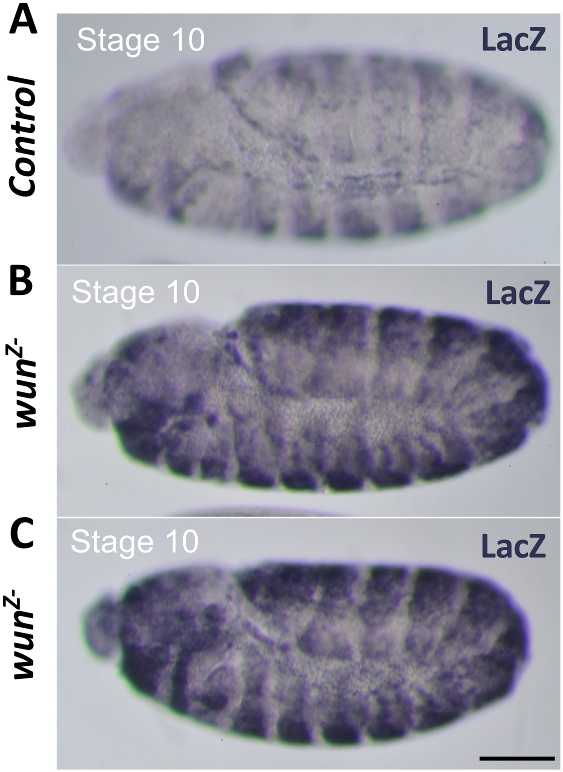


**Supplementary Figure 1.** **(A-C)** Recombinant *ptc-gal4 UAS-β-gal* females were mated with the *UAS-RNAi* males of the indicated genotype. Stage 10/11 embryos (F1) derived from the respective crosses were stained with anti-*β-gal* antibody. Compared to the *control, ptc-gal4 UAS-β-gal* (A), all embryos *ptc-gal4 UAS-β-gal/ UAS-RNAi* (n~30) show considerably enhanced β-*gal* expression, which indicates Ptc activation (B and C). Panel B & C are biological replicates.

Figure S2

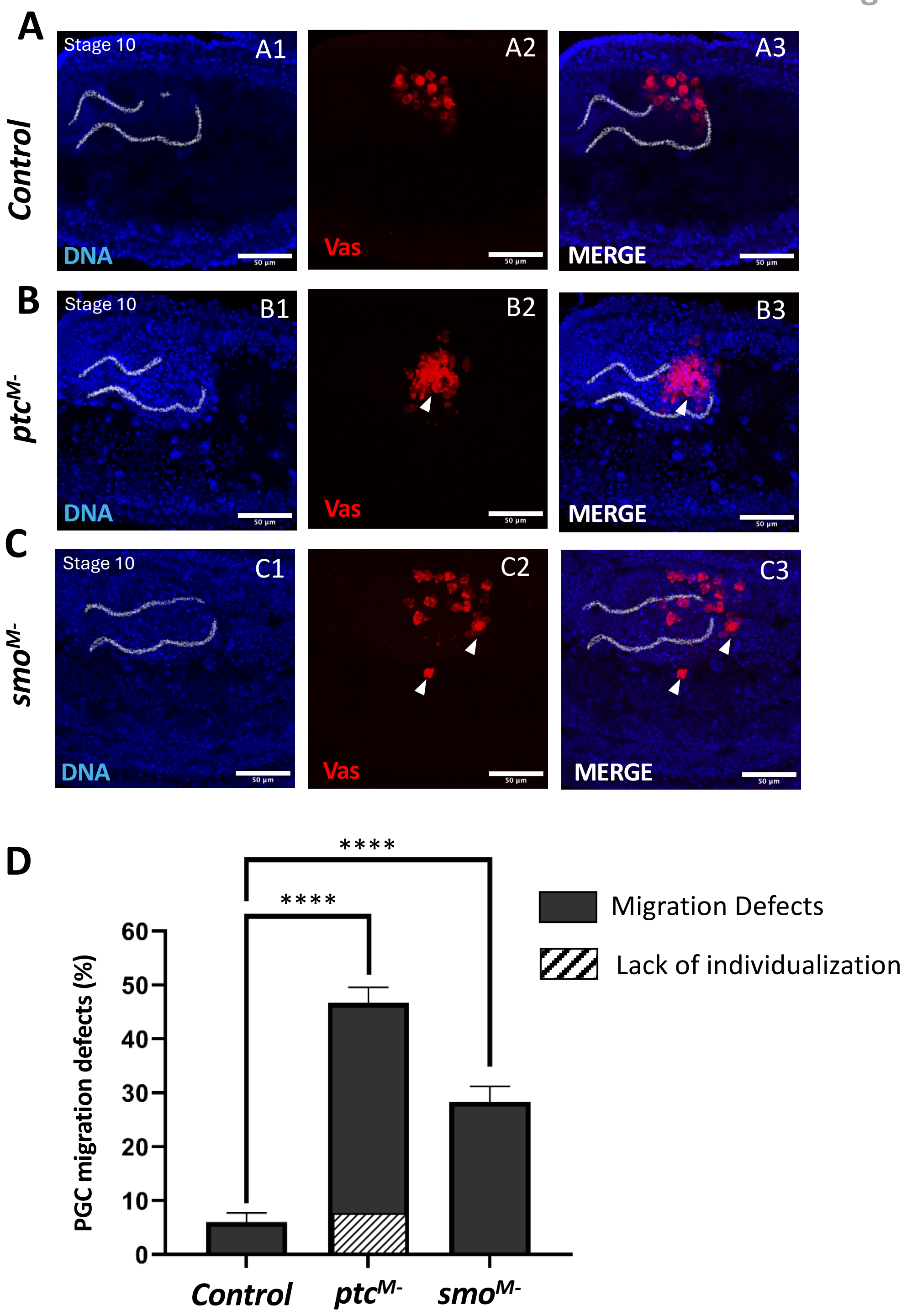


<u>Supplementary Figure 2</u>. **Maternal requirement of Hh receptors, *ptc*, and *smo* during PGC migration in early embryos.** (A-C) Stage 10 embryos maternally deprived of indicated genes were laid by the nos-gal4>UAS-RNAi of indicated genes. PGCs in control *lexA*$^{M-}$ embryo display clearly individualized PGCs that traverse across the gut through its dorsal side (A). In contrast, PGCs in *ptc*$^{M-}$ embryos either clump and fail to individualize altogether (B) or show delayed migration through the gut possibly due to incomplete individualization. Conversely, PGCs of *smo*$^{M-}$ (C) are able to individualize like the control PGCs and exit the gut but start to scatter on their way to the mesoderm. . Panels (A1-C1) display Hoechst staining, Panels (A3-C3) displayed merged images. Arrowheads mark mis-migrated embryos. Gut is outlined. (D) Aberrant migration of PGCs is quantified in graph using Ordinary one-way ANOVA and Tukey's multiple comparison test (n=12,3 replicates), ****p<0.0001. Diagonal lines represent percentage of embryos that have PGCs that have not individualized by stage10.

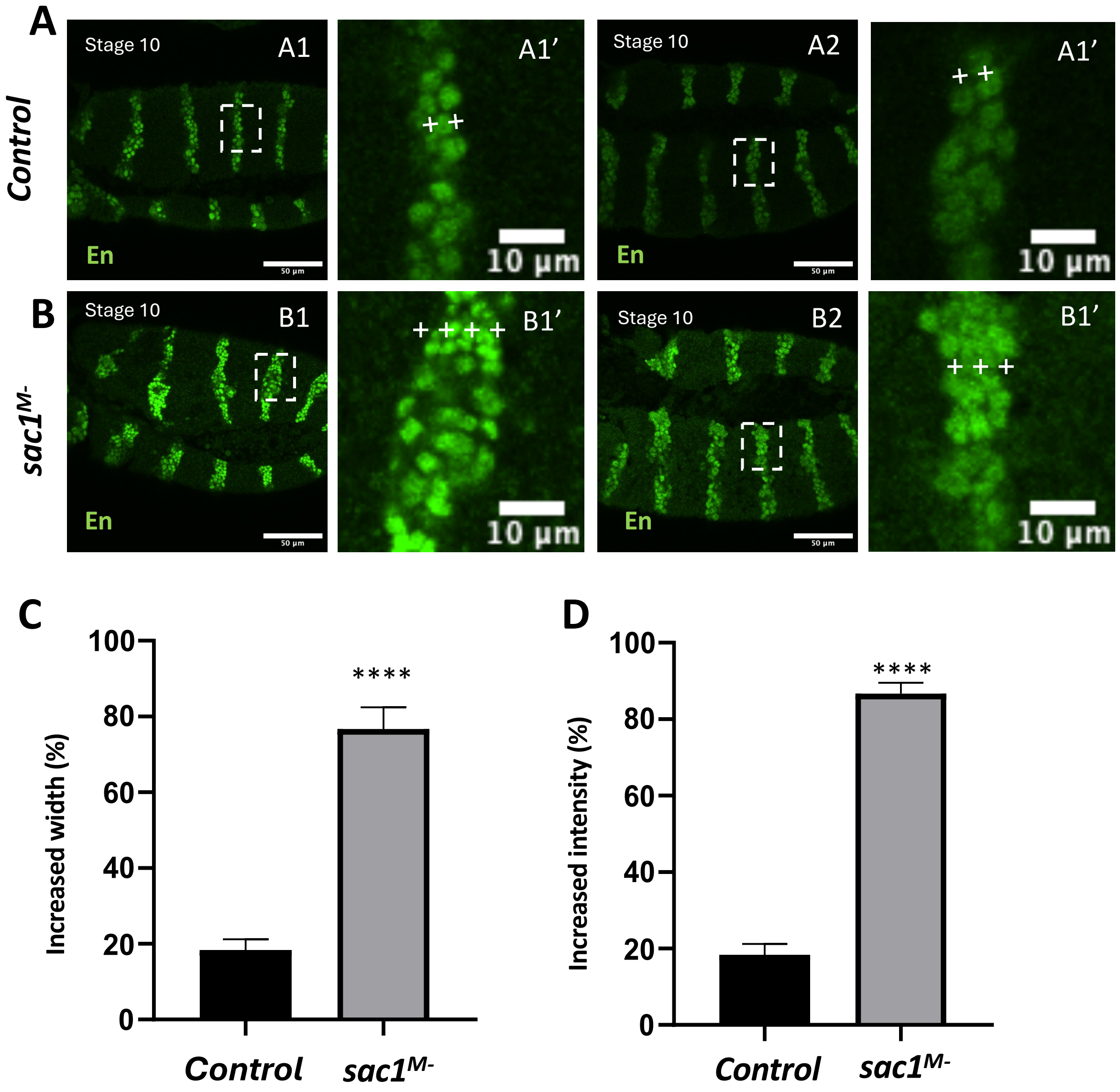


**Supplementary Figure 3. Embryos derived from mothers compromised for *sac1* (*sac1$^{M-}$*) show elevated En expression in the ectoderm.** (A-B) Stage 10 Embryos derived from *nos-gal4>UAS-RNAi* were stained with anti-Engrailed (En) antibody (green) and DNA dye, Hoechst (not shown*). Compared to the negative control *lex$^{M-}$* (A1 and A2) embryos a significantly higher percentage of *sac1$^{M-}$* (B1 and B2) embryos display expansion in the width of the individual stripes and increased level of En protein and. ‘+’ mark is used for the cells positive for En in a stripe. Panels (A1’,-A2’ and B1’- B2’) show magnified insets. (C-D) Quantification of En stripe broadening (C) and increased En expression (D) in embryos. Significance calculated using Ordinary one-way ANOVA (n=10 N=3), ****p<0.0001.

**Figure S4**

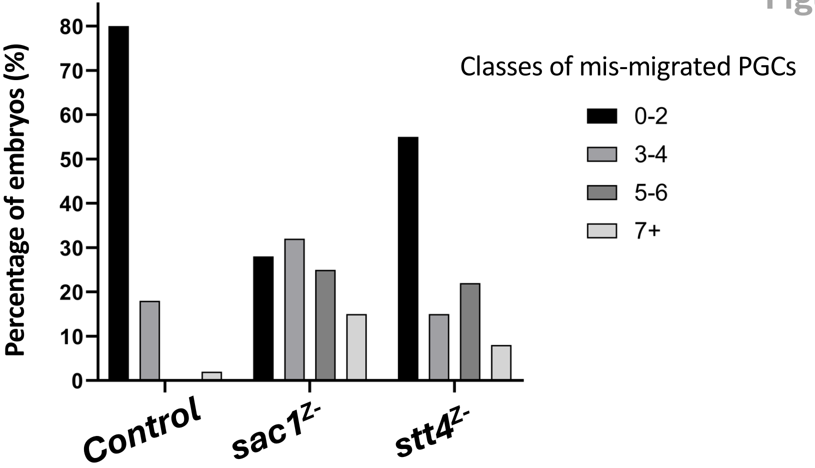


<u>Supplementary Figure 4</u>**. Zygotic depletion of lipid modification enzymes required for regulating Hh signalling results in germ cell migration defects**. Bar graph showing the percentages of embryos of indicated genotypes with a given number of mis-migrating germ cells. Chi-square Goodness of Fit test was done for each genotype with the *Control* (n = 38), $sac1^{Z-}$(n = 48) p = $2.01 \times 10^{-54}$, $stt4^{Z-}$ (n = 32) p = $4.83 \times 10^{-23}$

Figure S5

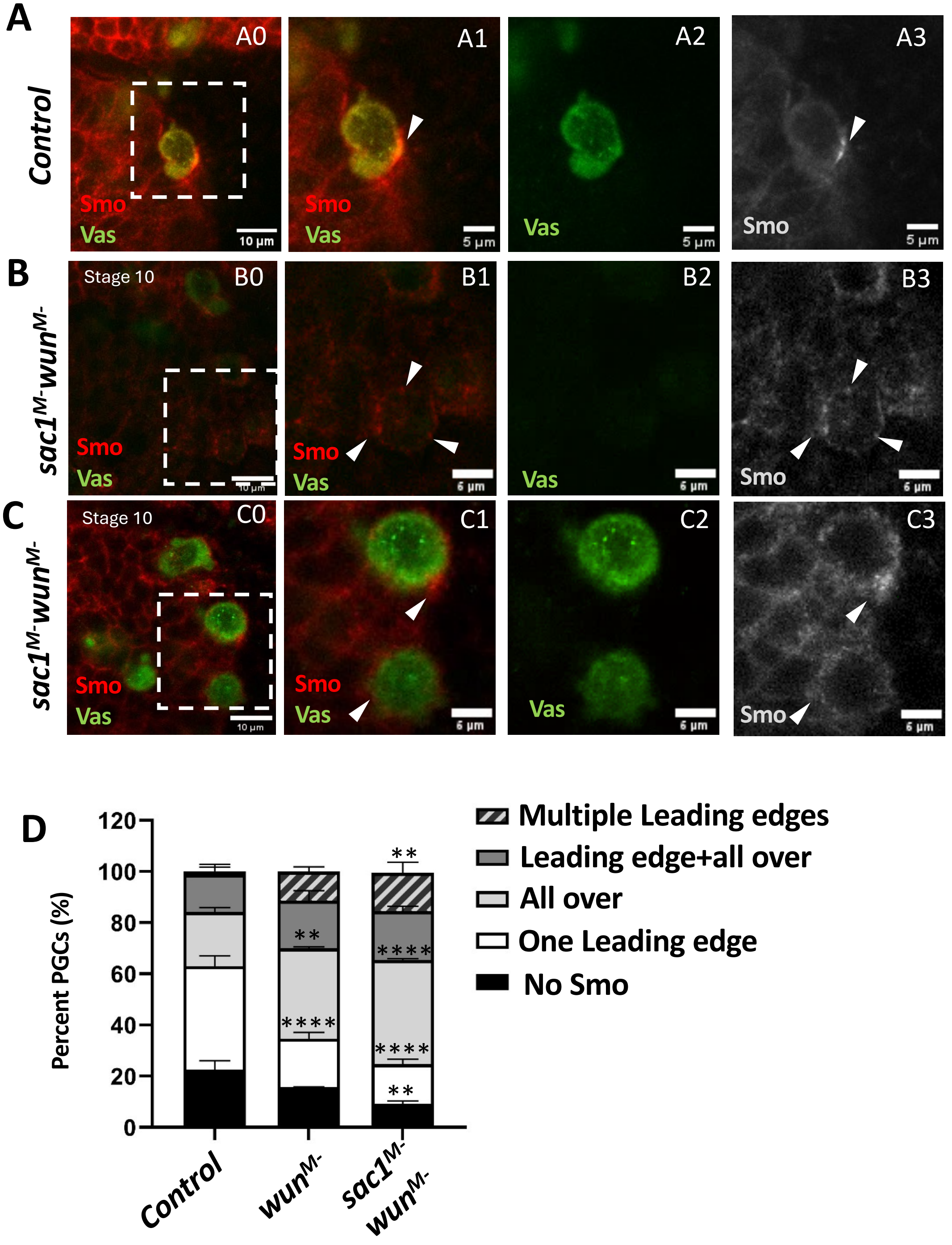


**Supplementary Figure 5. Genetic interaction between *wun* and *sac1* enhances aberrant Smo localization.** (A-C) Stage 10 embryos of the indicated genotype. In control *lexA*$^{M-}$embryos (A) Smo is localized at the leading edge (A0, magnified inset A1-A3). In contrast, in *sac1*$^{M-}$; *wun*$^{M-}$ PGCs, multiple foci of Smo localisation are observed (B0, magnified inset B1-B3) or are present all over the PGC (C0, magnified inset C1-C3). All panels were equally edited with minimum intensity value at 10 and maximum intensify value at 100 using Fiji Image J. (D) Comparison of percentages of embryos of indicated genotypes having Smo localization as per mentioned classes were quantified (n=40-70 across replicates) using 2-way ANOVA and Tukey's multiple comparison test *** p<0.001** p<0.01.
